# Immune-Derived THBS1-CD47 Axis Induces Cellular Senescence and Suppresses Osteogenesis in Diabetic Periosteum

**DOI:** 10.1101/2025.11.19.689158

**Authors:** Fangyuan Shen, Jianpeng Zhou, Moyu Liu, Qiaoyue Ren, Jingyao Cui, Puying Yang, Ting Zheng, Jun Wang, Ling Ye, Yu Shi

**Affiliations:** State Key Laboratory of Oral Diseases and National Clinical Research Center for Oral Diseases, West China Hospital of Stomatology, Sichuan University, Chengdu, China; Department of Endodontics, West China Hospital of Stomatology, Sichuan University, Chengdu, China

**Author notes:** Corresponding author: Yu Shi, Mailing address: State Key Laboratory of Oral Diseases & National Clinical Research Center for Oral Diseases, West China Hospital of Stomatology, Sichuan University, Chengdu, 610041, China;., Ling Ye, Mailing address: Department of Endodontics, West China Hospital of Stomatology, Sichuan University, Chengdu, China. These authors contributed equally to this work (Fangyuan Shen, Jianpeng Zhou).

**Keywords:** type 2 diabetes (T2DM), scRNA-seq, periosteum, osteogenesis

## Abstract

Type 2 diabetes mellitus (T2DM) weakens bone repair and increases fracture risk, but how the periosteal environment contributes to this problem is not fully understood. We found that diabetic mice showed thinning of the periosteum, reduced bone mass, and poor function of bone-forming cells. Using single-cell RNA sequencing, we discovered that immune cells and bone-lineage cells interact more strongly in diabetes through the THBS1-CD47 pathway. THBS1 was mainly produced by macrophages, while its receptor CD47 was abundant in osteogenic cells. Laboratory experiments showed that THBS1 blocked bone gene activity, increased inflammation and cell death, and damaged mitochondria, leading to higher oxidative stress. Knocking down CD47 reversed these effects, restoring bone cell activity and energy balance. In diabetic fracture models, blocking THBS1 improved callus formation and bone healing. These findings identify THBS1-CD47 as a key driver of periosteal dysfunction in T2DM and highlight it as a potential target to improve skeletal repair in diabetic patients.

## Introduction

T2DM is an increasingly prevalent global health concern, affecting over 400 million individuals worldwide (1). In addition to the well-recognized comorbidities, including cardiovascular disease, neuropathy, and kidney dysfunction, accumulating evidence demonstrates that T2DM is a significant risk factor for skeletal vulnerability and increased susceptibility to fractures (2, 3). Reduced bone quality and turnover, along with increased cortical porosity, contribute to the elevated fracture risk in individuals with diabetes. Diabetic bone disease is also exacerbated by chronic hyperglycemia, oxidative stress, and low-grade inflammation, which compromise osteoblast function, delay fracture healing, and promote bone marrow adiposity (4). Cellular senescence is a key contributor to diabetic bone fragility; senescent osteoblasts and mesenchymal stem cells exhibit reduced proliferative capacity, enhanced inflammatory cytokine secretion, and impaired osteogenic differentiation (5, 6).

The periosteum, a highly vascularized connective tissue layer surrounding bones, is crucial to skeletal homeostasis, adaptation, and fracture healing (7, 8). Bone injury reactivates periosteal progenitor, which contribute to soft and bony callus formation to support bone regeneration. Periosteal progenitor cells contribute more substantially to bone repair than bone marrow mesenchymal stromal cells (BMSCs) (9, 10). They also exhibit higher osteogenic and chondrogenic potential compared to BMSCs (11). Following injury, periosteal progenitors become reactivated and promote the formation of cartilage and bone callus, which serve as the main sources of bone regeneration. Although BMSCs play a critical role in bone homeostasis, diabetes-induced metabolic and osteogenic dysfunction of BMSCs has been well documented (12, 13). Moreover, periosteal cells are highly sensitive to microenvironmental changes, and chronic inflammation and aging-related dysfunction can disrupt osteogenic differentiation and impair regenerative capacity (14–16). Diabetes can cause persistent low-grade inflammation, leading to excessive immune cell infiltration, especially macrophages, which alter the periosteal niche and interfere with normal osteogenesis (17). Senescent cells accumulate in diabetic and aged bones and secrete senescence-associated secretory phenotype (SASP) factors including pro-inflammatory cytokines, chemokines, and proteases that contribute to chronic inflammation and bone fragility (18, 19). Pro-inflammatory cytokines, such as IL-17, IL-1β, and TNF-α, suppress osteoblast differentiation and promote osteoclastogenesis and bone resorption, further weakening bone integrity (20). This pro-inflammatory signal has been shown to increase oxidative stress and metabolic dysfunction by suppressing mitochondrial respiration and ATP production, leading to ROS elevation, cellular energy depletion, and disruption of osteogenic potential (21, 22). The accumulation of SASP factors in bone amplifies inflammatory signaling, reinforcing a non-regenerative, pro-fibrotic microenvironment that hinders fracture healing.

THBS1 is a multifunctional extracellular matrix glycoprotein that modulates cell adhesion, angiogenesis, and immune regulation, via CD47 and latent TGF-β activation (23). CD47 is a widely expressed transmembrane receptor that functions as a “don’t eat me” signal. CD47 prevents immune-mediated clearance of apoptotic cells by interacting with SIRPα on macrophages (24). This mechanism helps maintain cell survival under normal conditions. However, during aging, the upregulation of CD47 in osteolineage cells leads to evasion of apoptosis, altered immune surveillance, and prolonged inflammatory signaling, exacerbating bone loss and impairing skeletal homeostasis (25). Studies in aged mice demonstrated that THBS1 knockout reverses many of the detrimental effects, leading to the upregulation of osteogenic pathways in bone marrow stromal cells and the downregulation of inflammatory genes (26). Thus, the absence of THBS1 may enhance osteogenesis and reduce inflammation, supporting the hypothesis that THBS1-CD47 signaling contributes to skeletal aging and diabetic bone fragility.

Current evidence suggests that the periosteum is a critical yet understudied niche in diabetic bone disease. While diabetes is known to impair bone quality and fracture healing, most studies have focused on bone marrow–derived progenitors, leaving the periosteal compartment largely unexplored. Given its structural location and enrichment of osteogenic progenitors, we hypothesized that T2DM disrupts the function of periosteal osteogenic lineage cells and disturbs the immune–osteogenic microenvironment, ultimately leading to reduced bone quality and impaired fracture repair. To test this hypothesis, we investigated changes in periosteal structure, cellular composition, and osteogenic potential under diabetic conditions. Using single-cell RNA sequencing (scRNA-seq), we characterized diabetes-induced alterations in immune cell populations and their regulatory effects on periosteal progenitors, and further examined the impact of these changes on mitochondrial function, cellular senescence, and osteogenic activity. Finally, we evaluated whether targeted blockade of pathological signaling could improve fracture healing in diabetic mice. Collectively, these findings provide novel insight into periosteal pathophysiology in diabetes and highlight potential therapeutic strategies for enhancing skeletal regeneration in T2DM.

## Results

### Characterization of the T2DM Model and Periosteal Morphology in Mice

To establish a classic T2DM mouse model, C57BL/6J mice were fed a high-fat diet (HFD) for 6 weeks, followed by a single 100 mg/kg intraperitoneal streptozotocin (STZ) injection. Fasting blood glucose levels were measured one week after STZ administration, and mice with glucose levels below 11.1 mmol/L were excluded from this study. (Figure 1A and S1A, B). The remaining mice were then maintained on HFD for an additional 3 months. To verify the successful establishment of the T2DM model, hyperglycemia and impaired glucose tolerance were confirmed using intraperitoneal glucose tolerance tests and insulin tolerance test (Figure S1C and D). We further assessed femur and tibia length and wet weight showed no significant differences between CTRL and T2DM mice (Figure S1E–G). Femur from diabetic and non-diabetic mice were analyzed using micro-computed tomography (micro-CT) (Figure 1B). Bone mass and cortical and trabecular bone mineral density (BMD) were significantly reduced in T2DM mice compared with controls (Ctrl) (Figure 1C and D). Cortical bone area and thickness decreased in T2DM mice, reflecting a loss of bone tissue in this critical load-bearing region (Figure 1C). Similarly, trabecular bone volume fraction (BV/TV) was reduced, indicating a decline in trabecular quality (Figure 1D).

**Figure 1.**
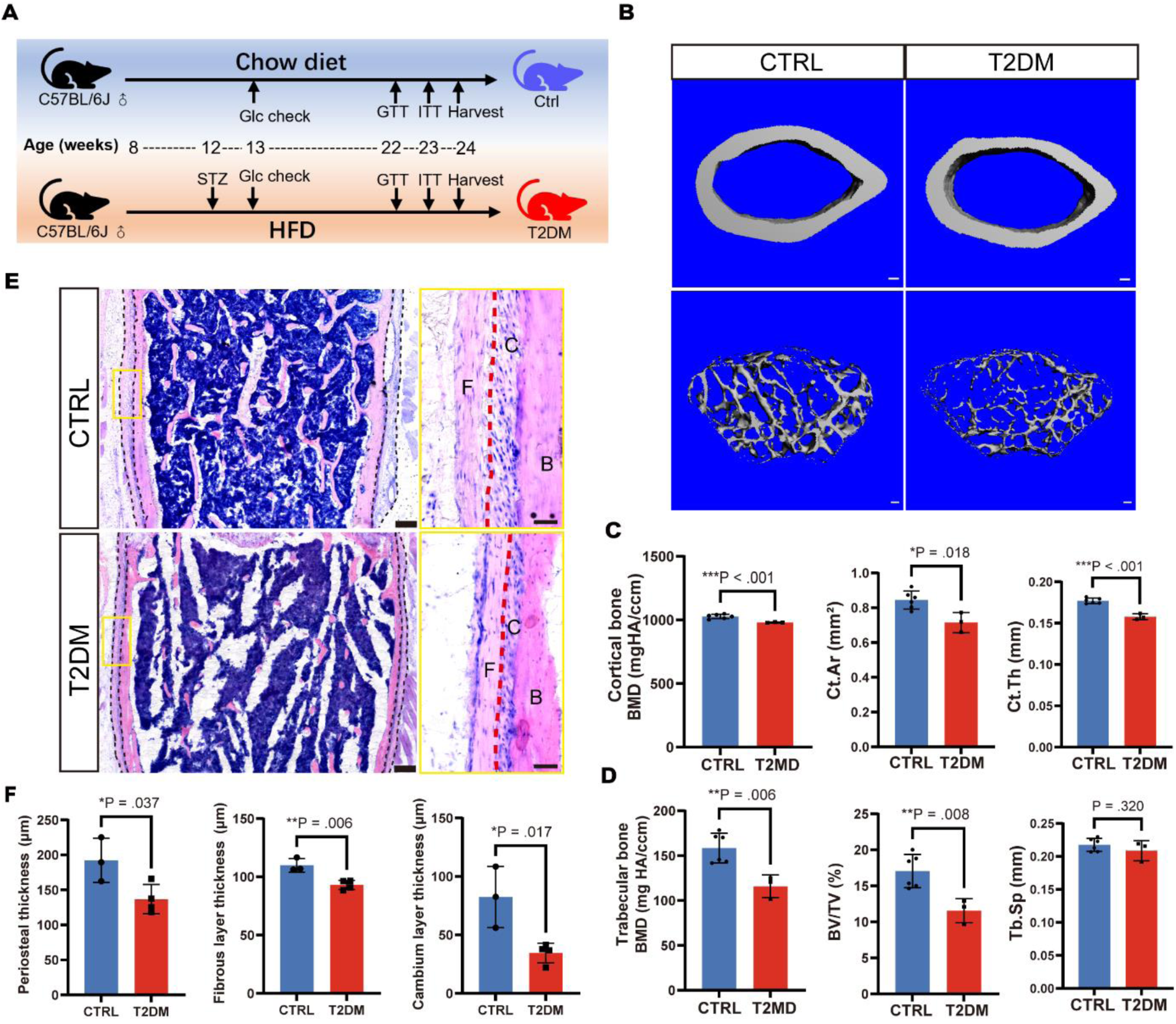
Bone and periosteal phenotypes in type 2 diabetic mice (**A**) A schematic for experimental design. STZ dose: 100 mg/kg body weight. (**B**) Representative µCT images. Scale bar: 100 μm. (**C**, **D**) Quantitative analysis of cortical bone (**C**) and trabecular bone (**D**) in femur. Biological replicates: CTRL, n=6 mice; T2DM, n=3 mice. (**E**) H&E staining of femoral bone. Boxed regions are presented at higher magnification to the right. The black dashed lines separated cortical bone and periosteum; the red dashed lines separated the fibrous layer and cambium of the periosteum. Thick scale bar: 100 μm, thin scale bar: 50 μm. (**F**) Quantification of periosteum area and thickness of fibrous and cambium layers. Biological replicates: CTRL, n=3 mice; T2DM, n=4 mice. All bar graphs are presented as mean ± S.D. Data were analyzed using unpaired student’s t-test with Welch’s correction. BMD, bone mineral density; Ct.Ar, cortical bone area; Ct.Th, cortical bone thickness; Tb.Sp, trabecular separation; F, fibrous layer; C, cambium layer; B, cortical bone.

The periosteum is essential for bone repair and maintenance. It comprises a fibrous layer for structural support and a cambium layer that serves as a niche for osteolineage cells, including skeletal stem cells, progenitor cells, and osteoblasts (27). In T2DM mice, the overall periosteal thickness was reduced, and the cambium and fibrous layers were significantly thinner (Figure 1E and F). The type 2 diabetic mice also exhibited significant bone damage, including reduced bone mass, lower cortical and trabecular BMD, and structural alterations in the periosteum. Diabetes-induced periosteal thinning likely contributes to impaired bone maintenance and repair, resulting in skeletal fragility.

### Impaired Proliferation and Survival of Periosteal Osteolineage Cells in T2DM

Postn-CreERT2 expression is a marker for periosteal mesenchymal progenitor cells, a subpopulation of cells critical for maintaining skeletal homeostasis and bone regeneration in vivo (28). Postn-CreERT2; tdTomato mice with established diabetes were treated with tamoxifen for five consecutive days after three months HFD feeding (Figure 2A and S1H).

**Figure 2.**
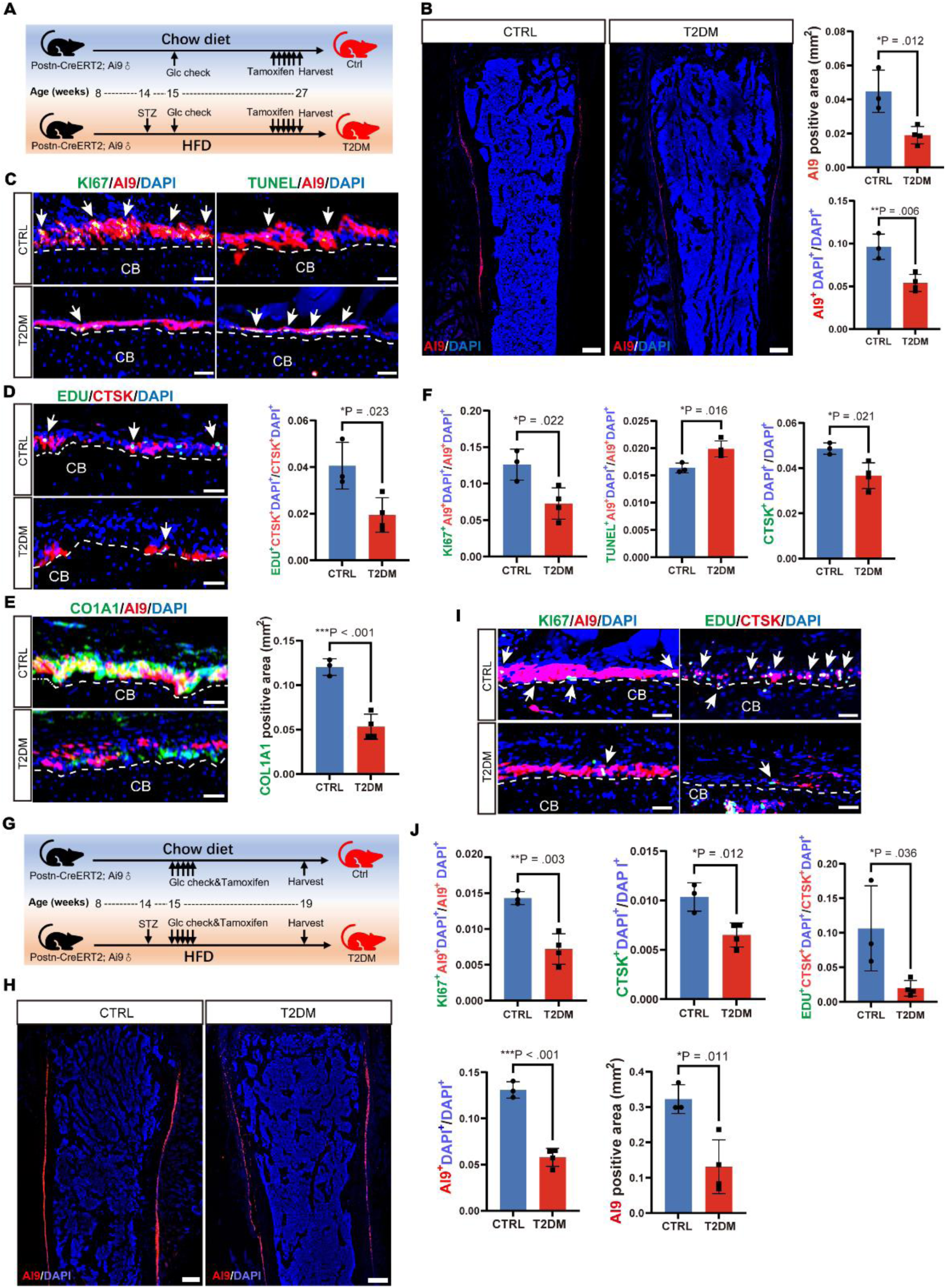
Dysfunction of POSTN+ periosteal progenitor cells in the periosteum of type 2 diabetic mice. (**A**) A schematic for experimental design. (**B**) Representative images and quantification of Postn reporter activity in the femur of Postn-CreERT2; tdTomato mice following a 5-day tamoxifen injection. Mice with 3-month diabetes (n=4) exhibited significantly reduced Postn-expressing periosteal cells compared to healthy controls (n=3). Scale bar: 200 μm. (**C-F**) Immunofluorescence staining and corresponding quantitative (**F**) bar graphs of KI67 & TUNEL (**C**) EdU & CTSK (**D**), COL1A1 (**E**) in the femur of diabetes and controls group. Scale bar: 50 μm. (**G**) Scheme depicting diabetes modeling and lineage tracing. (**H**) Distribution and quantification (lower right) of tdTomato+ cells in the femur of Postn-CreERT2; tdTomato mice 1 month after tamoxifen injection in diabetic (n=4) and healthy (n=3) group. Scale bar: 200 μm. (**I, J**) Quantification and immunofluorescence staining of KI67 and EdU in the femur of diabetes and controls group. Arrows indicate representative examples of fluorescent signal colocalization. Scale bar: 50 μm. All quantitative data are presented as mean ± S.D. Each data point represents one mouse (biological replicate). Statistical analysis was performed using unpaired Student’s t-test with Welch’s correction. CB, cortical bone.

Postn reporter activity was markedly reduced in T2DM group compared with Ctrl group (Figure 2B). Based on Ki67 staining of Postn-positive cells, proliferating periosteal progenitor cells were significantly reduced in diabetic mice (Figure 2C and F). CTSK is another periosteal stem cell marker (8). CTSK-positive cells and CTSK-positive proliferating cells were reduced in T2DM mice compared with Ctrl mice (Figure 2D and F). These results collectively demonstrate that T2DM weakens the proliferation ability of periosteal progenitor cells, potentially impairing bone repair and maintenance. Apoptosis of Postn-positive periosteal progenitor cells was significantly increased in T2DM mice, as evidenced by TUNEL staining (Figure 2C and F). In addition, staining for the osteogenesis marker COL1A1 was reduced in the periosteum of diabetic mice, reflecting impaired osteogenic activity (Figure 2E).

To investigate the effects of T2DM on periosteal lineage cells, tamoxifen was administered for five consecutive days to both Postn-CreERT2; tdTomato diabetic mice and Ctrl mice, followed by one month of continued high-fat diet or chow diet before sacrifice (Figure 2G and S1H). Immunofluorescence analysis revealed a decrease in tdTomato-positive cells (Figure 2H), and reduced co-localization with Ki67 (Figure 2I and J), indicating that the proliferation of Postn-positive periosteal lineage cells was impaired in the diabetic mice. CTSK staining showed a relative reduction in CTSK-positive cells in T2DM mice. EdU labeling confirmed that the proliferation of CTSK-positive cells was significantly reduced one month after the oneset of diabetes (Figure 2I and J). Overall, these findings suggest that T2DM impairs skeletal maintenance and repair by disrupting the balance between periosteal progenitor cell survival, proliferation, and osteogenic differentiation.

### Single-Cell Transcriptomic Analysis of Periosteum in T2DM

ScRNA-seq of the periosteum from T2DM and control mice was conducted to identify the molecular mechanisms underlying diabetes-induced defects. Integrated dataset analysis identified nine distinct cell populations based on marker gene expression (Figure 3A). Osteolineage cells (*Col1a1*, *Prrx1*, and *Pdgfra*) (29, 30) represented the primary bone-forming population, while neutrophils (*Retnlg*, *Ltf*, and *Ly6g*) (31) and macrophages (*Adgre1*, *Csf1r*, and *Cd68*) (32) were key components of the immune microenvironment (Figure 3B). Additional cell populations included myeloid progenitor cells, pro-B cells, mature B cells, T cells, mast cells, and dendritic cells. Based on cell ratio analyses, the proportion of osteolineage cells decreased and macrophages increased in the periosteum of diabetic mice, indicating that immune regulation in the periosteal microenvironment was altered (Figure 3C-E).

**Figure 3.**
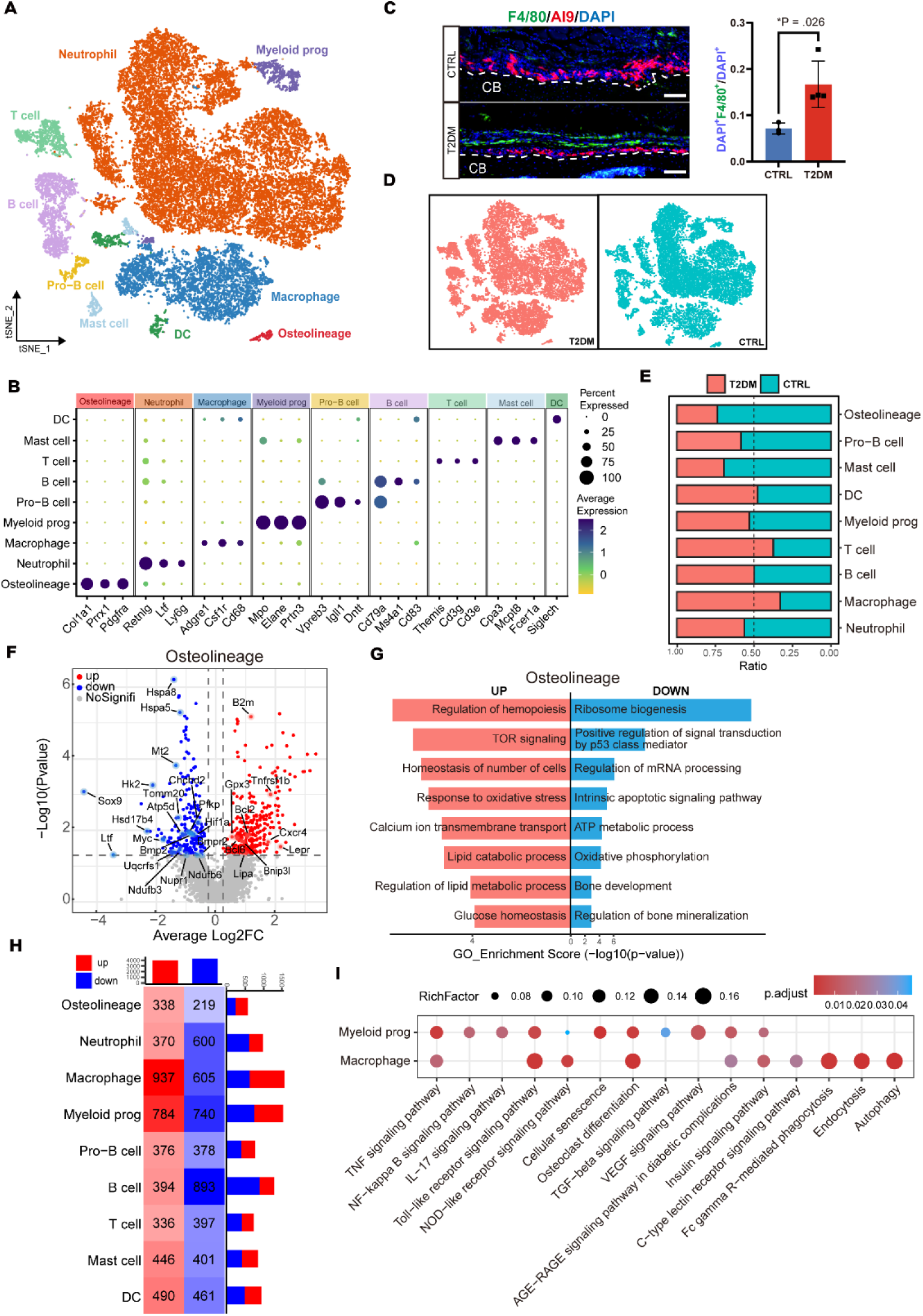
Single-cell transcriptomic analysis of the periosteum in diabetic and healthy mice. (**A**) tSNE plot depicting scRNA-seq clustering of integrated periosteal cells from T2DM and CTRL groups. (**B**) Dot plots of marker genes are used to annotate each identified cluster. (**C**) Immunofluorescence staining of periosteal macrophages (F4/80^+^) in Postn-CreERT2; tdTomato mice with quantification (right, mean ± S.D) of macrophage abundance. Biological replicates: CTRL, n=3 mice; T2DM, n=4 mice. Scale bar: 100 μm. Unpaired Student’s t-tests. (**D**) tSNE plots displaying separate distributions of periosteal cells from T2DM and CTRL groups, highlighting differences in cluster composition between conditions. (**E**) Bar graph showing the relative proportions of each cell cluster in T2DM and CTRL periosteal samples. (**F**) Volcano plot illustrating differentially expressed genes in osteolineage cells between diabetic and CTRL samples, with upregulated genes in red and downregulated genes in blue. (**G**) Gene Ontology (GO) enrichment analysis of differentially expressed genes in osteolineage cells, showing key biological processes affected by diabetes. (**H**) Heatmap of differentially expressed genes in each cluster between T2DM and CTRL, with red indicating upregulated genes and blue indicating downregulated genes. (**I**) KEGG pathway enrichment analysis of upregulated genes in myeloid progenitor cells and macrophages.

Transcriptional changes were detected in the osteolineage cells from T2DM mice. *Bmp2*, *Bmpr2*, and *Sox9*, which are crucial for osteoblast differentiation, extracellular matrix organization, and bone mineralization, were downregulated, suggesting that osteogenesis was impaired (Figure 3F). Glycolysis-related genes (*Hk2*, *Pfkp*, *Hif1a*, *Myc*, and *Nupr1*) and oxidative phosphorylation-related genes (*Chchd2*, *Ndufb3*, *Ndufa6*, and *Uqcrfs1*) were also downregulated, indicating mitochondrial dysfunction and reduced energy production. Gene enrichment analysis revealed that pathways related to bone development, oxidative phosphorylation, and ribosome biogenesis, which are essential for cellular function and osteogenic activity, were suppressed. Conversely, autophagy and fatty acid metabolism were upregulated, suggesting that the diabetic microenvironment induces metabolic adaptation. The ATP metabolic process and oxidative phosphorylation pathway were also enriched in T2DM mice, suggesting mitochondrial defects (Figure 3G). Autophagy-related genes, including *Bnip3l*, *Bcl6*, and *Bcl2*, were upregulated, indicating a complex regulatory balance between autophagy activation and inhibition (Figure 3F). The increased expression of *Bnip3l* is consistent with enhanced mitophagy (33), but *Bcl2* may counteract the initiation of autophagy by inhibiting Beclin-1 (34). In addition, the upregulation of fatty acid metabolism-related genes suggests altered energy utilization under diabetic conditions (Figure S2B).

Diabetes-induced changes were also evident in immune cell populations, including macrophages, myeloid progenitors, B cells, and neutrophils (Figure 3H). Immune regulation pathways, including TNF, IL-17, and Toll-like receptor signaling, were upregulated in myeloid progenitor cells and macrophages (Figure 3I). These pathways contribute to inflammatory responses and bone resorption (35, 36). Notably, osteoclast differentiation pathways were enriched in myeloid progenitors and macrophages, indicating a potential role in promoting osteoclast activity and accelerating bone loss in diabetes. Upregulated B cell receptor, T cell receptor, and NF-kappa B signaling pathways in B cells suggest enhanced immune activation, which may contribute to chronic inflammation in the periosteum. The proportion of neutrophils did not change significantly. However, genes related to lysosome function, autophagy, and sphingolipid metabolism were upregulated in neutrophils, indicating altered immune cell function and lipid processing in the diabetic microenvironment (Figure S2A and B). The upregulation of AGE-RAGE signaling in macrophages, neutrophils, and myeloid progenitor cells highlights a common response to hyperglycemia-induced oxidative stress that which may contribute to impaired bone homeostasis (Figure 3I).

Increased inflammatory activity in the periosteal microenvironment, driven by macrophage expansion and upregulation of pro-inflammatory pathways, may disrupts osteoblast function. The enhanced osteoclastogenic potential of myeloid progenitors and macrophages suggests an imbalance favoring bone resorption over formation. Metabolic dysfunction, including impaired mitochondrial activity and disrupted energy metabolism, also occurred in osteolineage cells, further compromising their capacity to maintain bone integrity. These findings highlight the mechanisms by which diabetes alters periosteal cell composition and signaling, leading to skeletal deterioration.

### Altered Cell Communication in Diabetic Periosteal and Bone marrow

A comparative analysis revealed that cell-cell communication networks in the periosteal niche were significantly altered in diabetic mice. Osteolineage cells exhibited enhanced regulation of immune cells, in diabetic samples, and the regulatory influence of myeloid progenitor cells and macrophages over osteolineage cells increased (Figure 4A and S2C). Thbs1-Cd47 interactions between neutrophils, macrophages, and bone lineage cells were upregulated, and Osm signaling was downregulated (Figure 4B, C and S2D), limiting osteogenic activation and bone formation (37). Metabolic dysfunction exacerbated these effects; downregulated mitochondrial and glycolysis-related genes impaired energy production, restricting osteolineage viability (Figure 3F). Expression analysis revealed increased *Thbs1* in macrophages and myeloid progenitors, and higher *Cd47* expression in osteolineage cells (Figure 4C and S2D). Flow cytometry demonstrated an increased proportion of CD47^hi^ periosteal cells in diabetic mice, indicating that cellular function was altered (Figure 4d and e). Immunofluorescence and ELISAs confirmed the increased THBS1 levels in the periosteal microenvironment in mice with diabetes (Figure 4F and G). Elevated *Cd47* expression impairs regenerative capacity and promotes dysfunction in aged muscle stem cells (38). This same phenomenon may occur in diabetes. CD47^hi^ increased in the periosteal cells of diabetic mice, indicating reduced self-renewal, increased susceptibility to apoptosis, and decreased periosteal repair capacity. This dysfunction likely exacerbates bone loss and metabolic imbalances, contributing to skeletal fragility in diabetes.

**Figure 4.**
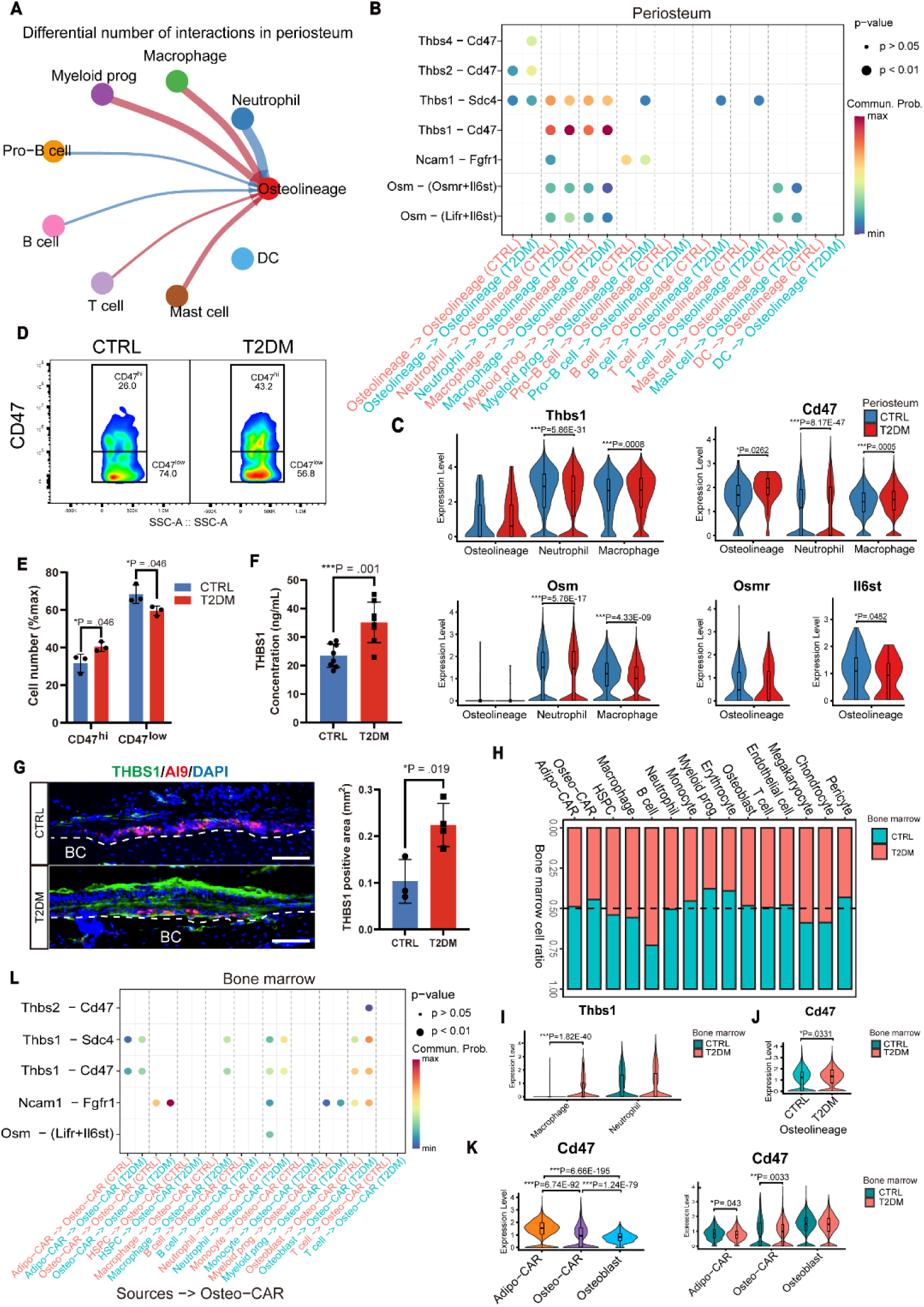
Cell-cell communication analysis in the periosteum and bone marrow of diabetic mice. (**A**) CellChat comparison analysis illustrating changes in the number of immune cells regulating osteolineage cells in the periosteum of diabetic and control group. Red lines indicate increased regulation, while blue lines indicate decreased regulation, with line thickness representing the interaction strength. (**B**) Differential ligand-receptor pairs between T2DM and CTRL periosteal cells. (**C**) Violin plots of periosteal expression levels of ligands and receptors involved in THBS1 and OSM signaling. Statistical significance was assessed using MAST. (**D**) Flow cytometry analysis measuring CD47 protein expression in periosteal cells from CTRL (n=3 mice) and 3-month T2DM (n=3 mice). (**E**) Bar graph comparing the relative abundance of CD47^hi^ and CD47^low^ periosteal cells in 3-month T2DM and CTRL mice. (**F**) ELISA quantification of THBS1 levels in serum from T2DM (n=8 mice) and CTRL (n=8 mice). (**G**) Immunofluorescence staining of THBS1 in periosteum, with quantification of fluorescence intensity. CTRL, n = 3 mice; T2DM, n = 4 mice. Scale bar: 100 μm. (**H**) Relative proportion of bone marrow T2DM and CTRL in each cell cluster. (**I**-**K**) Expression levels of Thbs1 and Cd47 in immune cells and osteolineage cells of bone marrow. Statistical significance was assessed using MAST. (**I**) Differential ligand-receptor interactions between immune cells and osteo-CAR cells in the bone marrow of T2DM and CTRL group. Bar graphs represent mean ± S.D. Statistical analysis was performed using unpaired Student’s t-test unless otherwise stated.

The scRNA-seq data from osteolineage (12) and immune cells (13) in the bone marrow were integrated to validate the changes in THBS1-CD47 signaling in diabetic bone (Figure S3A and B). Using previously annotated cell types and marker genes, 15 distinct cell types were identified, but no significant alterations in the overall composition of bone marrow mesenchymal cells were detected (Figure 4H and S3C-E). However, a slight reduction in the Osteo-CXCL12 abundant reticular (CAR) cell population was observed in the diabetic group. The proportions of B cells and macrophages increased in diabetic samples, indicating a potential shift toward a more inflammatory bone marrow microenvironment (Figure 4H). This result is consistent with previous studies demonstrating that immune cell dynamics, are altered and may contribute to impaired bone regeneration in diabetes (13). *Thbs1* levels in macrophages, and the proportion of *CD47*-expressing cells in the Osteo-CAR cluster increased in bone marrow of the diabetic group, indicating a shift in osteolineage regulation (Figure 4I-K and S3F). *Cd47* expression differed among the osteolineage cells. Cd47 was expressed at the highest expression levels in Adipo-CAR cells, followed by Osteo-CAR cells; osteoblasts exhibited the lowest expression of Cd47 (Figure 4K). Interaction analysis revealed that THBS1-CD47 regulation of the Osteo-CAR, Adipo-CAR, and osteoblasts by macrophages and neutrophils significantly increased in the diabetic bone marrow environment, mirroring the changes observed in the periosteum (Figure 4L and Figure S3G, H).

To further dissect immune alterations in periosteum versus bone marrow, we performed flow cytometry to quantify macrophages and neutrophils under CTRL and T2DM conditions. The results demonstrated that macrophage proportions were significantly increased in both periosteum and bone marrow of T2DM mice, whereas neutrophil abundance remained unchanged (Figure 5A, B). qPCR analysis of sorted immune populations revealed compartment-specific patterns of Thbs1 expression. In T2DM mice, Thbs1 was significantly upregulated in macrophages from both periosteum and bone marrow. Notably, periosteal macrophages exhibited consistently higher Thbs1 expression levels compared to their bone marrow counterparts, underscoring their role as a major local source of THBS1 in the diabetic bone niche (Figure 5C). These findings suggest that periosteal macrophage-derived THBS1 drives a shift toward enhanced inflammatory and stress-induced signaling, thereby impairing the regenerative capacity of osteolineage cells. The strengthened THBS1–CD47 interactions may promote cellular senescence, oxidative stress, and apoptosis, particularly in osteo-primed stem/progenitor cells essential for bone formation, ultimately reinforcing inflammatory signaling pathways that disrupt osteogenesis and contribute to skeletal fragility in T2DM.

**Figure 5.**
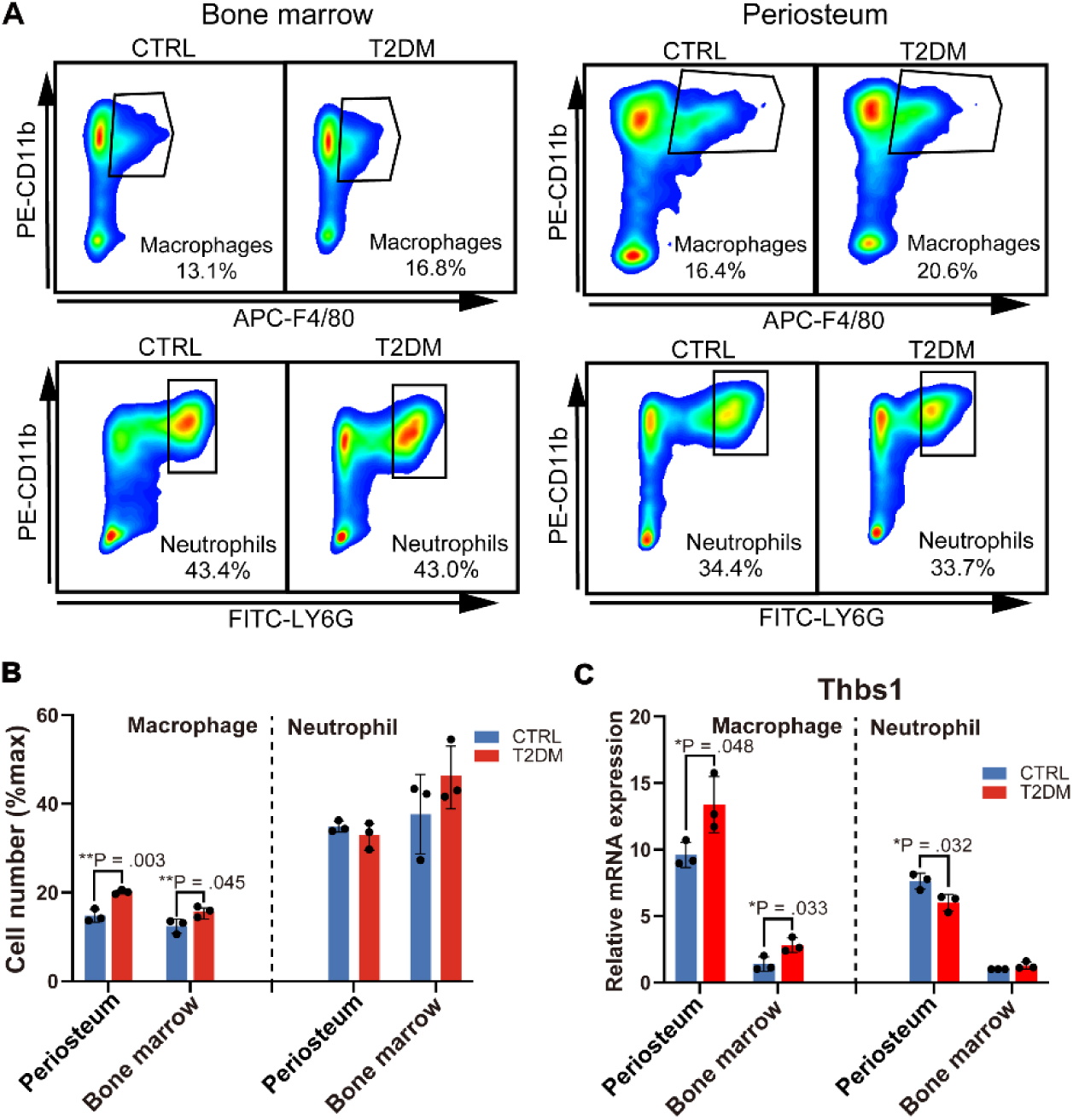
Flow cytometry and qPCR analysis of macrophages and neutrophils in periosteum and bone marrow. (**A**) Representative flow cytometry plots showing gating of macrophages and neutrophils from bone marrow and periosteum in CTRL and T2DM mice (n=3 mice/group). (**B**) Quantification of the proportions of macrophages and neutrophils in periosteum and bone marrow. (**C**) Relative mRNA expression of Thbs1 in flow-sorted macrophages and neutrophils from periosteum and bone marrow as determined by qPCR. Data are presented as mean ± S.D. Statistical analysis was performed using unpaired Student’s t-test.

### THBS1 Impairs Periosteal Cell Function and Fracture Healing in Diabetes

THBS1 is a secreted protein that regulates cell behavior via interactions with extracellular matrix components and cell surface receptors. It plays pivotal roles in regulating angiogenesis (39), wound healing (40), inflammation (41), and cancer (42). Accumulation of THBS1 in the extracellular matrix has been associated with aging (43) and aging-related chronic diseases, including obesity (44), fatty liver disease (45), and diabetes (46). In vitro experiments were conducted to investigate the effects of THBS1 on periosteal cell function. Periosteal cells were treated different concentrations of recombinant THBS1, and EdU proliferation assays and migration tests were used to determine the threshold concentration. THBS1 significantly reduced the proportion of EdU+ proliferating cells and inhibited cell migration, suggesting that THBS1 inhibits periosteal cell renewal and motility (Figure 6A-D). The increase in senescent cells after THBS1 treatment was confirmed with senescence-associated β-galactosidase (SA-β-gal) staining (Figure 6E). These results indicate that THBS1 directly promotes periosteal cell aging.

**Figure 6.**
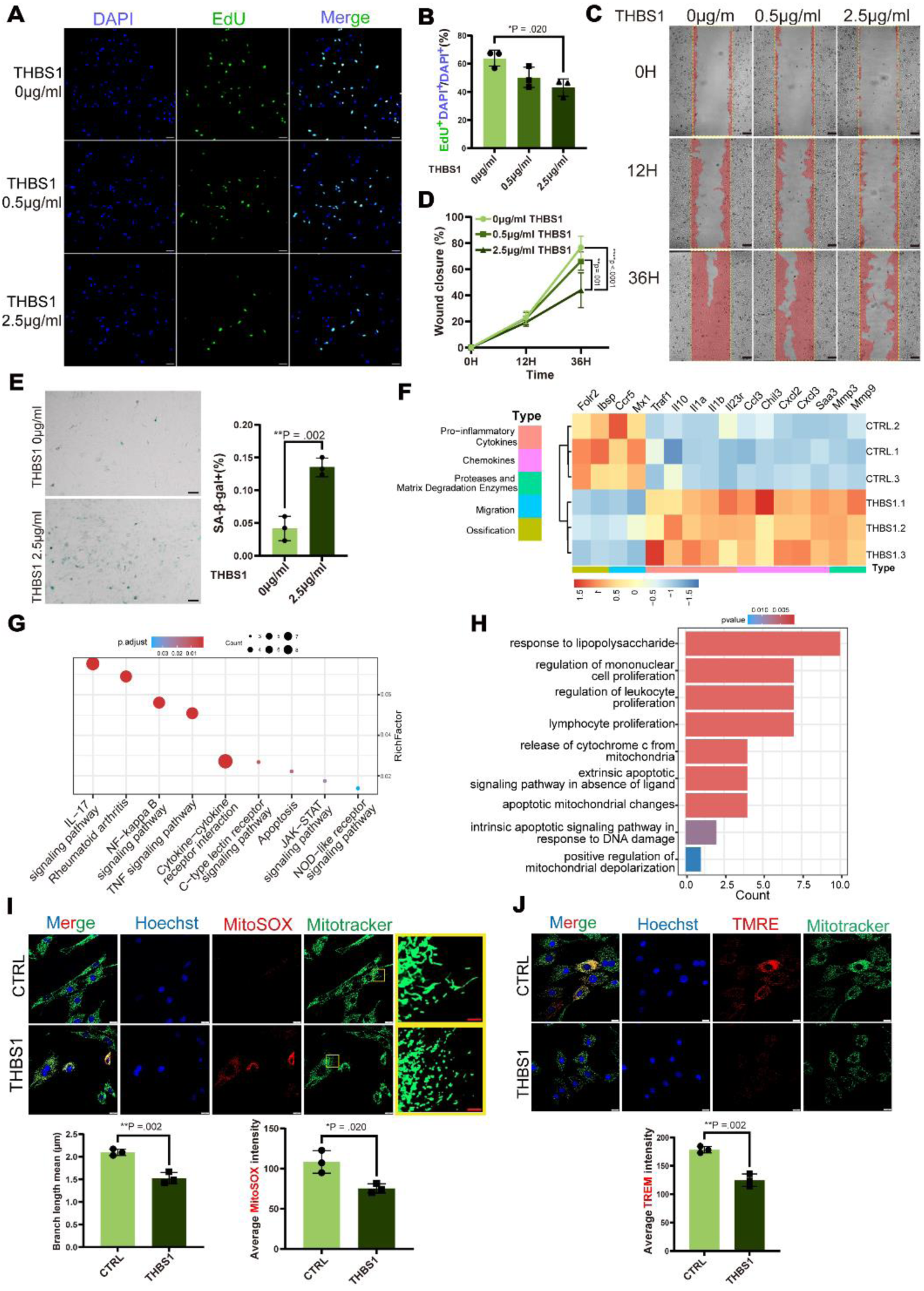
THBS1 impairs periosteal cell proliferation, migration, and promotes cellular senescence. (**A**, **B**) Representative images of EdU cell proliferation assay (**A**) and quantification (**B**) of periosteal cells three days after different concentrations of THBS1 treatment. Scale bar: 50μm. (**C**, **D**) Periosteal cell migration assay (**C**) and quantification (**D**) of relative wound closure percentage under different concentrations of THBS1 treatment. Scale bar: 100μm. (**E**) Representative SA-β-gal staining images and percentages (right) of senescent cells in periosteal cells in response to THBS1 treatment. Scale bar: 100μm (**F**) Heatmap of differentially expressed genes from RNA-seq analysis comparing THBS1-treated periosteal cells (n = 3 biological replicates) and controls (n = 3 biological replicates). (**G, H**) KEGG pathway (**G**) and GO term (**H**) enrichment analysis of upregulated DEGs in THBS1-treated periosteal cells, highlighting the activation of inflammatory, apoptotic, and mitochondria dysregulation pathways. (**I, J**) Representative fluorescence images and quantification of mitochondrial morphology in periosteal cells stained with MitoTracker, mitochondrial superoxide levels using MitoSox probe (**I**), and assessment of mitochondrial membrane potential using TMER dye (**J**). For mitochondrial morphology and functional assays, three biological replicates were performed, 24 randomly selected cells per group were analyzed. White scale bar: 50 μm, red scale bar: 5 μm. All data are presented as mean ± S.D. Biological replicates, n = 3 per group. Unpaired student’s t-test (two groups) or one-way ANOVA followed by Tukey’s post hoc test (multiple groups).

RNA-seq of THBS1-treated periosteal cells was employed to clarify the molecular changes induced by THBS1. THBS1 downregulated genes related to osteogenesis and migration (10), and upregulated pro-inflammatory cytokines, chemokines, and matrix metalloproteinases, all of which are key components of SASP (Figure 6F). Enrichment analysis revealed significant upregulation of pathways associated with inflammation and apoptosis, including IL-17 signaling, NF-kappa B, TNF, and JAK-STAT. (Figure 6G). These findings suggest that THBS1 suppresses osteogenic function and induces an inflammatory, senescence-like state in periosteal cells, impairing their regenerative capacity. Apoptosis-related mitochondrial changes were also enriched in THBS1-treated periosteal cells (Figure 6H). Mechanistically, THBS1 binding to CD47 mediates the translocation of DRP1 from the cytoplasm to the mitochondria (47, 48). DRP1 activation disrupts the mitochondrial electron transport chain, leading to mitochondrial membrane depolarization and increased ROS production (48). We examined the effects of THBS1 on periosteal cell mitochondria. Mitochondrial staining revealed a morphological shift from a continuous filamentous network to a fragmented, punctate structure, indicative of mitochondrial dysfunction (Figure 6I). In addition, mitochondrial ROS levels and membrane potential were evaluated using live-cell mitochondrial-targeted fluorescent dyes. MitoSOX Red fluorescence increased in THBS1-treated cells, indicating elevated mtROS production. Tetramethylrhodamine ethyl ester (TMRE) staining was reduced in THBS1-treated cells, indicating mitochondrial depolarization and impaired function (Figure 6I and J). These findings demonstrate that THBS1 induces an inflammatory, senescence-like state in periosteal cells and disrupts mitochondrial integrity, potentially exacerbating cellular dysfunction and impairing regenerative capacity.

To clarify the temporal sequence of THBS1-induced cellular dysfunction, periosteal cells were treated with THBS1 for 0, 12, 24, and 72 h, with or without the mitochondrial ROS scavenger Mito-TEMPO (Mito-T). SA-β-gal staining revealed a significant increase in senescent cells only at 72 h (Figure 7A and B), whereas mitochondrial alterations occurred earlier. Specifically, MitoSOX intensity increased as early as 12 h, accompanied by a decline in mitochondrial membrane potential and shortened mitochondrial branch length by 24 h (Figure 7D-F). These changes were largely rescued by Mito-T, which normalized ROS levels, restored membrane potential, and preserved mitochondrial morphology. Consistently, qPCR analysis demonstrated that the expression of senescence markers p16 and p53 was significantly upregulated at 72 h in the THBS1 group but attenuated by Mito-T (Figure 7I and J). Together, these results suggest that mitochondrial dysfunction and ROS accumulation precede the induction of cellular senescence, indicating a causal role of ROS in THBS1-mediated periosteal progenitor impairment.

**Figure 7.**
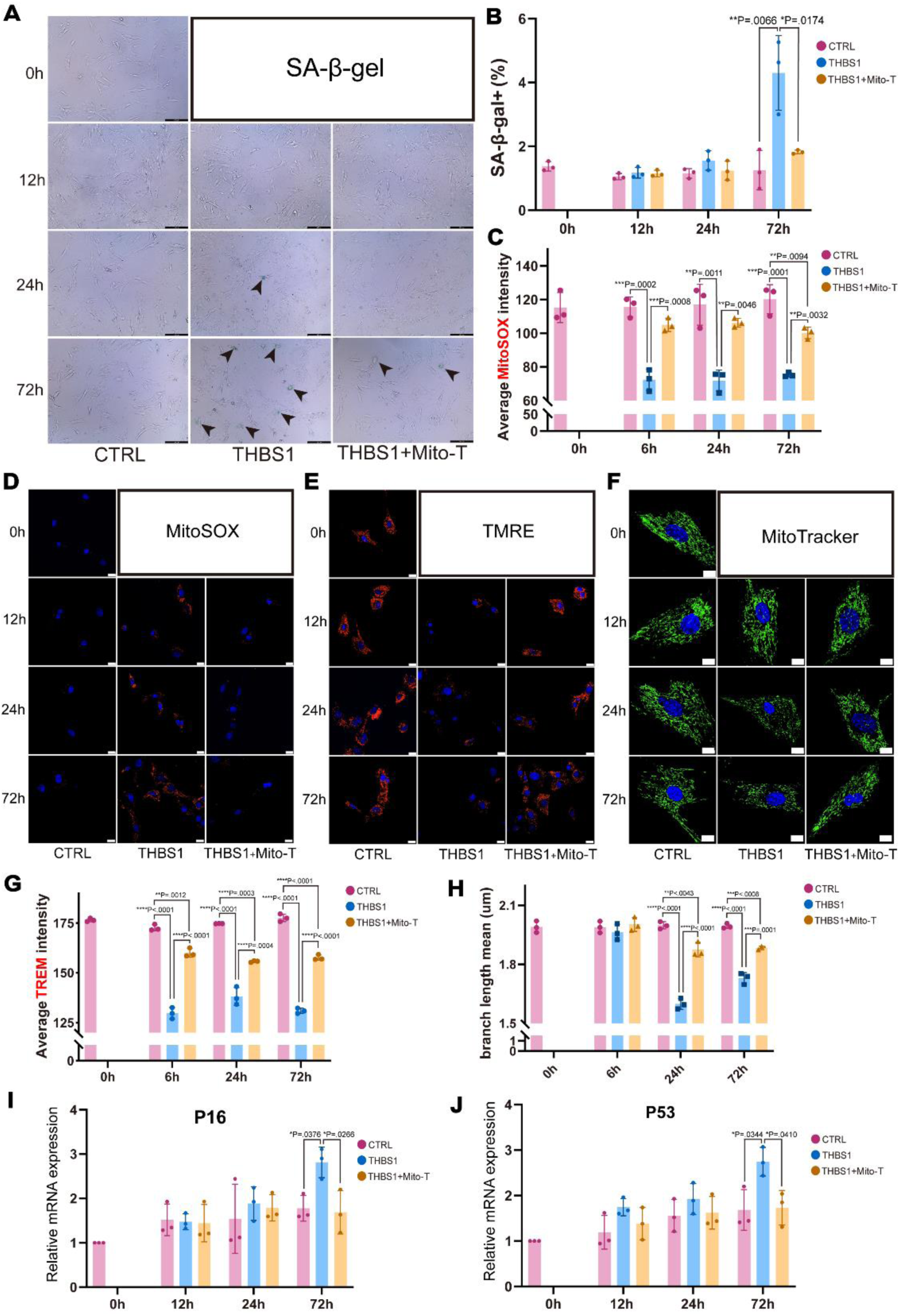
Time-course analysis of THBS1-induced mitochondrial dysfunction and senescence in periosteal cells. (**A**, **B**) Representative images (**A**) and quantification (**B**) of SA-β-gal–positive senescent cells under CTRL, THBS1, and THBS1 + Mito-T conditions at 0, 12, 24, and 72 h. Scale bar: 200 μm. (**C**) Quantification of MitoSOX fluorescence intensity at indicated time points. (**D**–**F**) Representative fluorescence images of MitoSOX (**D**), TMRE (**E**), and MitoTracker (**F**) staining in each group. Scale bars: 20 μm (thin), 10 μm (thick). (**G**–**H**) Quantification of TMRE intensity (**G**) and mitochondrial branch length (**H**). (**I**, **J**) Relative mRNA expression of p16 (**I**) and p53 (**J**) at indicated time points. Data are presented as mean ± S.D. For mitochondrial morphology and functional assays, three biological replicates were performed, 24 randomly selected cells per group were analyzed. For SA-β-gal and qPCR assays, three independent biological replicates were used. Statistical analysis was performed using one-way ANOVA followed by Tukey’s multiple comparisons test.

Since the periosteum contains the critical progenitors taking charge of bone fracture healing, we next assessed the impact of THBS1 on diabetic fracture healing. A mouse fracture model was established three months after the induction of T2DM (Figure S4A). Mice were locally injected with THBS1-blocking antibodies or an IgG control after the fracture (Figure 8A). THBS1 blockade significantly improved fracture healing in diabetic mice compared with the IgG control. Of note, micro-CT analysis revealed that bone density, bone volume fraction, and trabecular bone separation in the callus were restored to Ctrl levels in mice receiving the THBS1-blocking antibodies (Figure 8B and C). Histological analysis demonstrated that THBS1 inhibition increased woven bone formation in the callus and prevented the delayed cartilage-to-bone transition associated with diabetic fracture healing (Figure 8D). These results support THBS1 as a potential therapeutic target for improving bone repair in individuals with diabetes.

**Figure 8.**
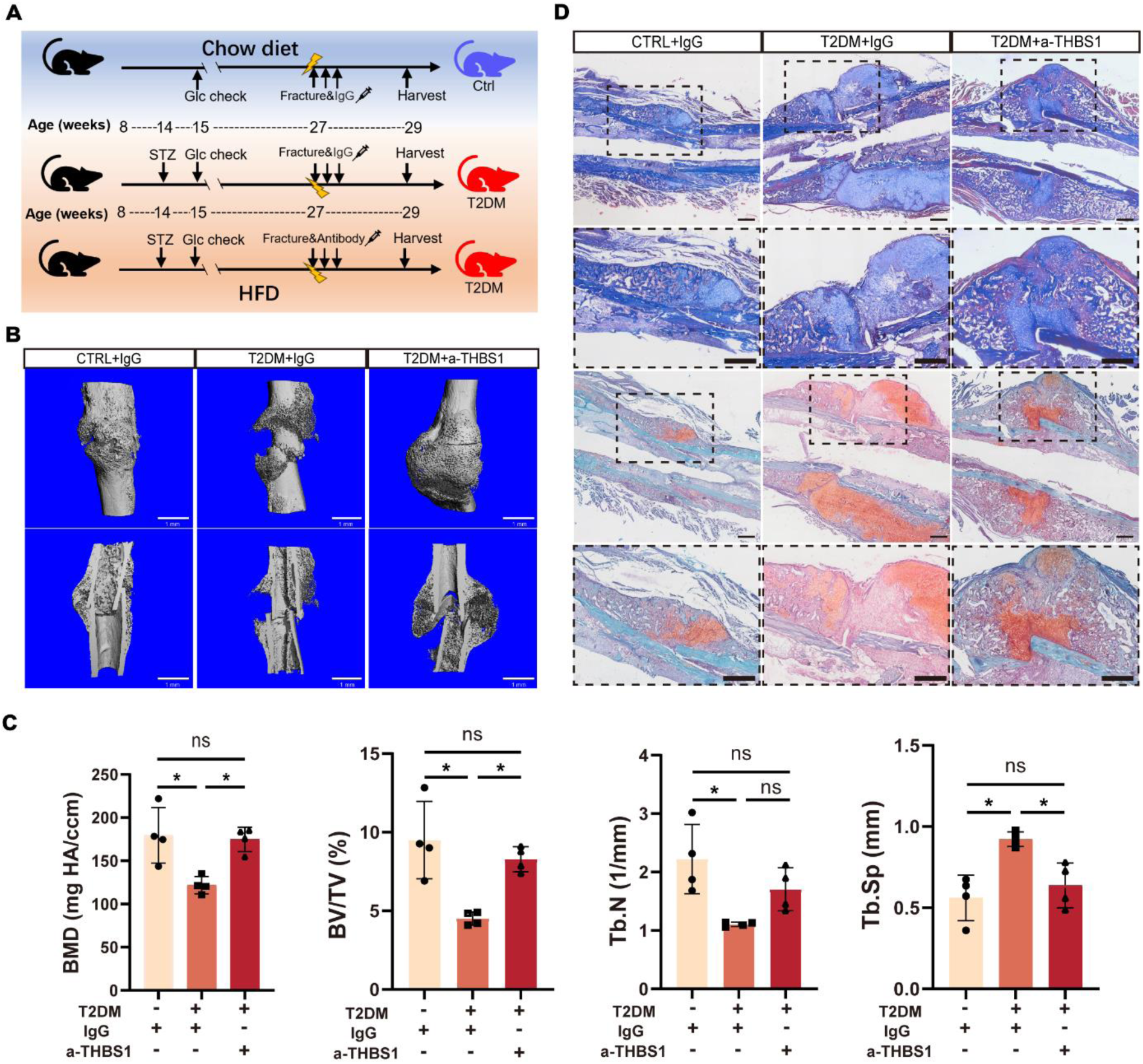
THBS1 blockade improves fracture healing in diabetic mice. (**A**) Schematic representation of the diabetic fracture model and post-fracture treatment groups. (**B**) Representative 3D micro-CT reconstruction images of femoral callus formation at day 14 post-fracture. Scale bar: 1 mm. (**C**) Quantitative micro-CT analysis of the femoral callus at day 14 post-fracture. (**D**) Representative histological images of Masson and Safranin O staining of medial femoral fracture sections at day 14 post-fracture, showing differences in cartilage and bone matrix composition across experimental groups. Higher magnification views of the areas outlined with dashed boxes. Thick and thin scale bar: 500 μm. Data are presented as mean ± S.D. Four biological replicates (n = 4 mice per group) were analyzed. Statistical comparisons were performed using one-way ANOVA followed by Tukey’s multiple comparisons test. *p<0.05; **p<0.01, ***p<0.001

### CD47 Knockdown Attenuates THBS1-Induced Senescence and Dysfunction in Periosteal Cells

To explore the role of CD47 in mediating THBS1-induced periosteal cell dysfunction, we examined the effects of THBS1 treatment after CD47 knockdown with shRNA in periosteal cells. The DNA damage marker, γ-H2AX staining, increased in THBS1-treated cells. However, this effect mitigated by CD47 knockdown. Thus, CD47 knockdown may protect against THBS1-induced genomic instability (Figure 9A). SA-β-gal staining revealed that THBS1 increased the number of senescent cells, and shCD47 significantly reduced senescence induced by THBS1, supporting the role of CD47 in THBS1-driven aging phenotypes (Figure 9B). Western blot analysis demonstrated that THBS1 reduced the expression of Lamin B1, a nuclear lamina protein whose loss is associated with cellular senescence (Figure 9C). In addition, qPCR analysis revealed that THBS1 treatment markedly upregulated the levels of critical genes associated with cellular senescence, including P16 and P53, whereas CD47 knockdown significantly attenuated their expression (Figure 9D). Notably, CD47 knockdown restored Lamin B1 protein levels, suggesting that Lamin B protects against THBS1-induced senescence.

**Figure 9.**
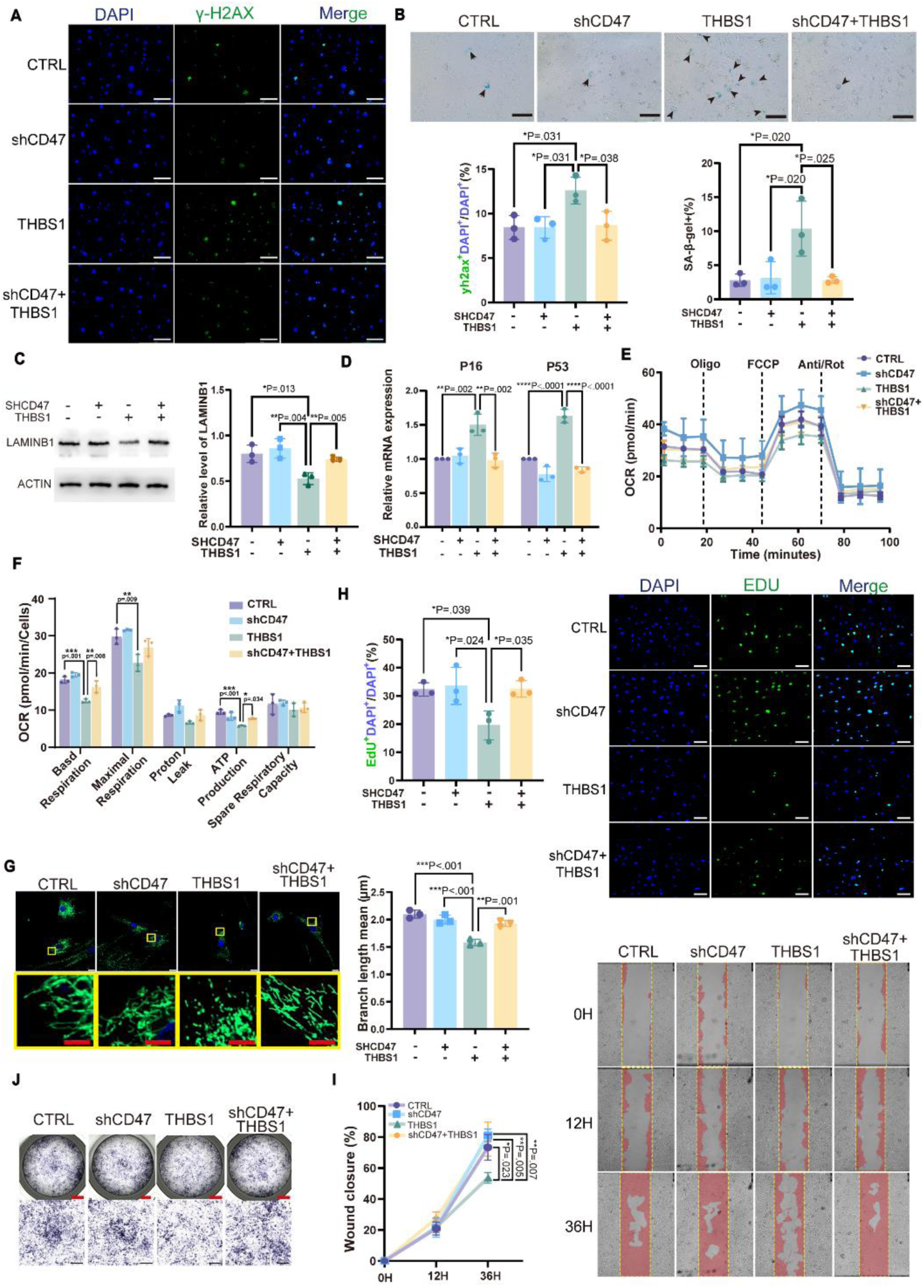
CD47 knockdown mitigates THBS1-induced cellular senescence and dysfunction in periosteal cells. (**A**) Representative images and quantification of γ-H2AX staining experiment detecting DNA damage, an indicator of cellular stress and senescence. Scale bar: 200 μm. (**B**) Representative images and quantification of SA-β-gal staining, measuring the proportion of senescent cells. Scale bar: 200 μm. (**C**) Western blots analysis of LAMINB1 are shown to evaluate senescence in periosteal cells. (**D**) Quantitative qPCR analysis of senescence-associated genes P16 and P53 expression. (**E**) Seahorse metabolic assay, assessing mitochondrial respiration and metabolic function across experimental groups. (**F**) Quantitative analysis of Seahorse assay parameters, comparing metabolic activity and energy production across groups. (**G**) Representative fluorescence images of MitoTracker. White scale bar: 50 μm, red scale bar: 5 μm. For each experimental group, 25 cells were randomly selected for mitochondrial quantitative analysis. (**H**) Representative images and quantification of EdU staining assessing periosteal cell proliferation across experimental groups. Scale bar: 200 μm (**I**) Representative images of migration assay evaluating the wound closure ability of periosteal cells in each group. Scale bar: 100 μm. (**J**) Representative images of ALP staining, assessing osteogenic differentiation potential among different conditions. Red scale bar: 2 mm, black scale bar: 1 mm. Data are presented as mean ± S.D. Three biological replicates (n = 3 independent experiments) were performed. For mitochondrial assays, three biological replicates were performed, 24 randomly selected cells per group were analyzed. Statistical analysis was performed using one-way ANOVA followed by Tukey’s multiple comparisons test.

Seahorse metabolic analysis was performed to assess the role of CD47 in THBS1-induced mitochondrial defects. CD47 knockdown alleviated THBS1-induced mitochondrial dysfunction in periosteal cells (Figure 9E). THBS1 treatment led to reduced oxygen consumption, compromised mitochondrial respiratory activity, and decreased ATP synthesis, indicating a decline in mitochondrial function and energy metabolism (Figure 9F). CD47 knockdown restored mitochondrial function, improving respiration and ATP production. MitoTracker staining showed that mitochondrial morphological changes from a normal reticular structure to discrete dots, indicating mitochondrial fragmentation and dysfunction, were induced by THBS1. CD47 knockdown prevented the reticular mitochondrial morphological changes induced by THBS1 (Figure 9G). Thus, THBS1 disrupts mitochondrial metabolism in periosteal cells, and CD47 knockdown mitigates these effects, highlighting CD47 as a key regulator of mitochondrial function and a potential therapeutic target for restoring metabolic homeostasis in diabetic bone regeneration.

THBS1 treatment significantly reduced EdU⁺ proliferating cells and impaired migration ability of periosteal cells, while CD47 knockdown attenuated these inhibitory effects (Figure 9H and I). Similarly, osteogenic differentiation assessed by ALP staining revealed that THBS1 suppressed ALP activity, which was restored upon CD47 knockdown (Figure 9J). To extend these findings in vivo, we locally injected a CD47-neutralizing antibody into fracture sites of T2DM mice (Figure 8A and S4B). Micro-CT analysis demonstrated that CD47 blockade markedly improved bone healing compared to IgG-treated T2DM mice, as evidenced by increased BMD, trabecular number, and bone volume fraction, alongside reduced trabecular separation (Figure 10A and B). Histological evaluation with Masson’s trichrome and Safranin O/Fast Green staining further confirmed enhanced callus formation and bone remodeling following CD47 antibody treatment (Figure 10C). Collectively, these results demonstrate that CD47 mediates THBS1-induced dysfunction of periosteal cells and that its inhibition rescues impaired osteogenesis and fracture repair under diabetic conditions.

**Figure 10.**
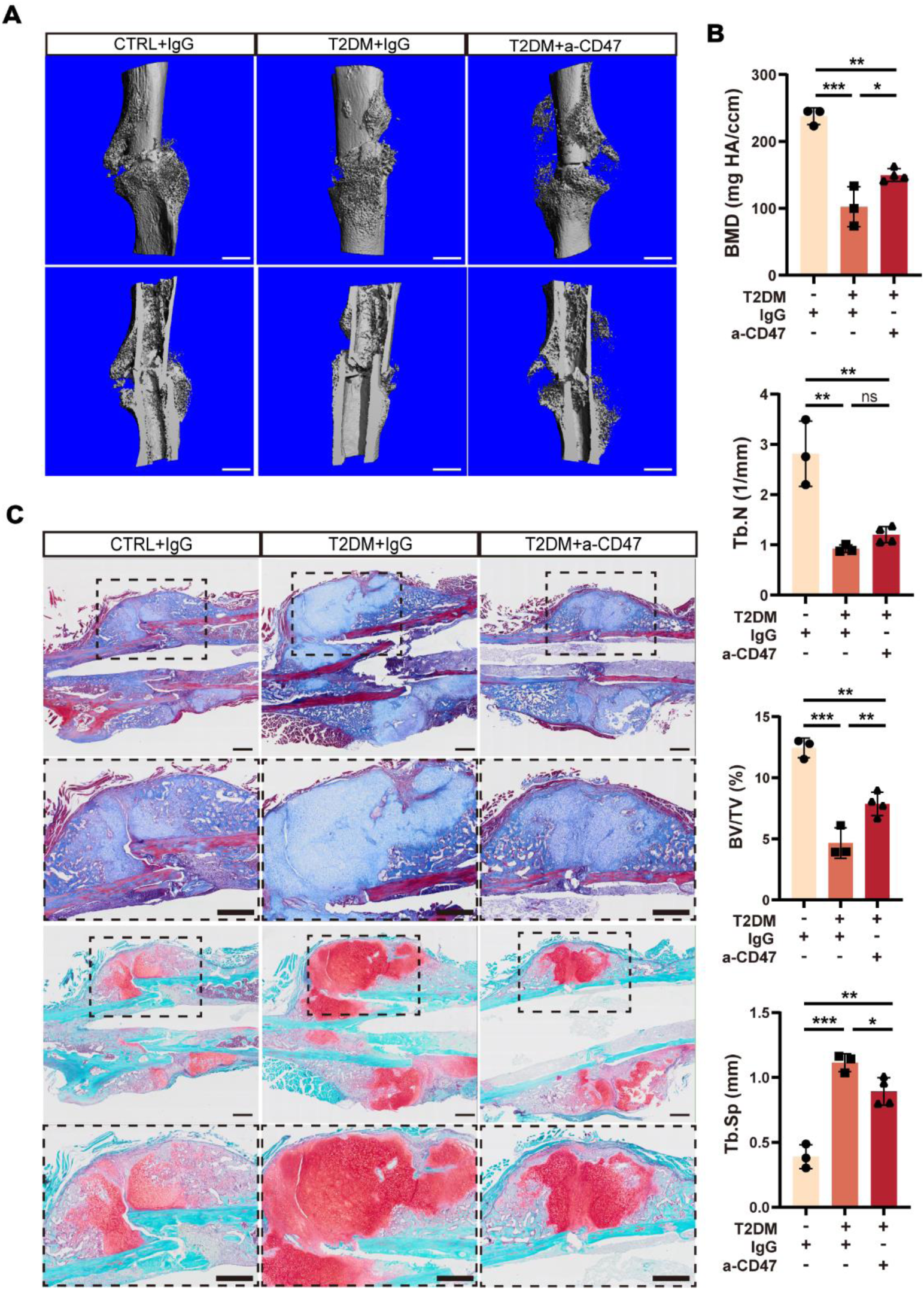
CD47 inhibition improves fracture healing in T2DM mice. (**A**) Representative 3D micro-CT reconstructions of femoral fracture callus formation at day 14 post-fracture in CTRL+IgG (n = 3), T2DM+IgG (n = 3), and T2DM + a-CD47 (n = 4) groups. Scale bar: 1 mm. (**B**) Quantitative micro-CT analysis of bone mineral density (BMD), trabecular number (Tb.N), bone volume fraction (BV/TV), and trabecular separation (Tb.Sp). (**C**) Representative histological staining of fracture callus with Masson’s trichrome and Safranin O/Fast Green, showing improved callus formation in T2DM+a-CD47 compared to T2DM+IgG. Higher magnification views of the areas outlined with dashed boxes. Thick and thin scale bar: 500 μm. Data are presented as mean ± S.D. At least three biological replicates were analyzed. Statistical significance was determined using one-way ANOVA followed by Tukey’s multiple comparisons test. *p<0.05; **p<0.01, ***p<0.001

## Discussion

Diabetic bone disease is characterized by altered bone metabolism, increased marrow adiposity, and osteoblast dysfunction, leading to reduced bone quality, progressive bone loss, impaired bone regeneration, and increased fracture risk. T2DM also causes chronic hyperglycemia, oxidative stress, and low-grade inflammation, which contribute to osteoblast dysfunction, mesenchymal stem cell depletion, and periosteal impairment (3, 49). T2DM is characterized by a chronic pro-inflammatory state with elevated cytokines, including TNF and IL-6. This chronic pro-inflammatory state promotes osteoclastogenesis and inhibits osteoblast differentiation, leading to impaired bone regeneration and an increased susceptibility to fracture. T2DM may also cause cellular senescence. Senescent osteoblasts secrete inflammatory and matrix-degrading factors that impair bone remodeling (2, 6). Thus, the immune microenvironment highly regulates the functions of osteolineage cells and the homeostasis of long bones. The detailed mechanisms of delayed bone regeneration after injury, in diabetes are unclear and worth addressing at a single-cell level.

Single-cell transcriptomic analysis of bone marrow–derived cells revealed high *Thbs1* expression in macrophages and a *Cd47* expression gradient in osteolineage cells: highest in Adipo-CAR cells, moderate in Osteo-CAR cells, and lowest in mature osteoblasts (Figure 4K). This pattern suggests that CD47 helps preserve progenitor cell identity by preventing immune clearance. In diabetic conditions, the proportion of cells expressing Cd47 increased in Osteo-CAR cells, while the expression of Cd47 in osteoblasts was downregulated. The increased *Cd47* expression in Osteo-CAR cells likely reflects a compensatory response to chronic oxidative and metabolic stress, promoting cellular resistance to immune-mediated phagocytosis. This compensatory response may help maintain progenitor cells; however, functionally impaired or senescent cells accumulate, resulting in a non-productive inflammatory niche. Conversely, the decreased *Cd47* expression may render mature osteoblasts more susceptible to immune-mediated clearance. This finding is consistent with previous studies focused on age-related bone loss. In aging individuals, Cd47 expression is downregulated in apoptotic osteoblasts, leading to preferential removal by macrophages, which eventually leads to osteoblast depletion and compromised bone formation (25).

THBS1 is a pro-inflammatory and pro-senescence cytokine. THBS1 promotes a senescent, inflammatory phenotype in periosteal mesenchymal progenitor cells and suppresses osteogenic differentiation. Our RNA-seq data showed that the increased expression of SASP-related cytokines and MMPs increased, consistent with the assumption that THBS1 drives tissue inflammation and age-related dysfunction (43, 50). Prior studies showed that THBS1-CD47 signaling inhibits mitochondrial respiration (51) and NO signaling (52, 53), increases oxidative stress (54, 55), and activates the p53/p21 axis to induce senescence (56, 57). In aged hematopoietic stem cells, THBS1 deficiency prolongs self-renewal and reduces inflammatory signaling (26). Furthermore, THBS1 levels are elevated in diabetic tissues and are associated with impaired regeneration (58). These data collectively demonstrate that THBS1-CD47 signaling remodels the diabetic periosteal niche into a non-regenerative, inflammatory state, disrupting osteolineage progression and impairing bone repair.

THBS1-CD47 signaling also drives senescence by promoting oxidative stress and mitochondrial dysfunction. THBS1 binding to CD47 inhibits nitric oxide signaling, leading to reduced mitochondrial respiration, elevated ROS, and ATP depletion (48). Consistent with these previous studies, THBS1 reduced basal and maximal respiration, diminished ATP production, and lowered spare respiratory capacity in periosteal cells. Furthermore, THBS1 increased mtROS and mitochondrial depolarization. These data demonstrate that THBS1-CD47 disrupts mitochondrial homeostasis and accelerates aging. Notably, the knockdown of CD47 rescued the mitochondrial impairments induced by THBS1, restoring energy metabolism and reducing senescence markers.

In addition to promoting senescence, THBS1-CD47 disrupts osteogenesis via interactions with the TGF-β1 pathway. Previous studies demonstrated that periosteal CD68⁺F4/80⁺ macrophages can promote bone formation through THBS1-mediated activation of latent TGF-β1 in response to mechanical loading (59). By contrast, the elevation of TGF-β1 levels in diabetes alters the physiological role of TGF-β1, shifting it from an osteo-inductive to an osteo-suppressive factor (60, 61). Xu et al. reported that excessive TGF-β1 impairs bone regeneration by altering Smad3 binding at the *Bmp2* promoter, to repress Bmp2 transcription and osteoblast differentiation (62). Our findings are consistent with this pathogenic shift. In the diabetic periosteum, increased THBS1 expression is primarily derived from macrophages, which may amplify TGF-β1 activation and reinforce a senescent and inflammatory microenvironment that suppresses osteogenic signaling. These results underscore the context-dependent role of THBS1–TGF-β1 signaling; under chronic metabolic stress, such as diabetes, THBS1–TGF-β1 signaling becomes maladaptive and contributes to osteolineage dysfunction and impaired bone regeneration. Macrophages act as the dominant source of THBS1 in the diabetic periosteal niche, reinforcing chronic inflammation and senescence. Normally, macrophages support bone remodeling and osteoblast function. However, diabetes skews their role toward inflammatory cytokine secretion and induction of senescence, thereby suppressing osteogenesis and contributing to skeletal fragility (18, 63). The upregulation of THBS1–CD47 interactions in macrophages further strengthens this inflammatory feedback loop, inhibiting osteoblast differentiation and impairing fracture healing. These findings identify THBS1–CD47 signaling as a key driver of diabetic bone fragility, where excessive TGF-β1 activation and macrophage-derived inflammation disrupt the balance between bone formation and degeneration.

Together, our findings delineate a multifaceted role for THBS1–CD47 signaling in diabetic bone disease, whereby immune-derived inflammation, mitochondrial stress, and impaired growth factor signaling converge to disrupt skeletal repair. Blockade of this pathway rescued osteolineage dysfunction and improved fracture healing in T2DM mice; however, translation requires careful consideration of safety and model limitations. CD47 is a central “don’t eat me” signal, and systemic inhibition carries risks of hematologic toxicity, immune dysregulation, and infection, while THBS1 is indispensable for host defense, as Thbs1–/–mice display impaired survival and heightened susceptibility to inflammatory injury (64, 65). Notably, although both THBS1 and CD47 blockade improved fracture healing, the effect of CD47 inhibition was less pronounced, which may reflect either intrinsic differences in their roles within the periosteal niche or variations in effective drug concentration. Our findings that local antibody delivery improved healing without organ toxicity suggest that targeted, localized intervention may represent a safer therapeutic strategy (Figure S5). At the same time, we acknowledge that the HFD/STZ-induced T2DM model, while recapitulating persistent hyperglycemia, impaired bone quality, and delayed fracture healing, does not fully reflect the chronicity and complexity of human T2DM. Thus, further studies are warranted to assess long-term safety, infection susceptibility, and validate these findings in models more closely mimicking human disease before clinical translation.

## Materials and Methods

### Experimental Design

The objectives investigated how T2DM impairs bone regeneration by disrupting the periosteal microenvironment. Using a HFD/STZ-induced T2DM mouse model, we assessed cortical bone structure and fracture healing by micro-CT and histology. Single-cell RNA sequencing profiled periosteal and bone marrow niches, revealing immune–osteogenic communication changes. Flow cytometry and qPCR validated Thbs1 expression in sorted immune populations. Functional assays tested THBS1 effects on periosteal progenitors, while CD47 knockdown and local antibody blockade were used to evaluate therapeutic rescue of osteogenic dysfunction and impaired fracture healing.

### Animals Samples

This study was approved by the Ethics Committees of West China School of Stomatology, Sichuan University (approval number WCHSIRB-AT-2025-380). To induce T2DM, eight-week-old male C57BL/6J mice were fed a high-fat diet (HFD; 60 kcal% fat) ad libitum. After six weeks on HFD, mice received a single intraperitoneal dose of streptozotocin (STZ; 100 mg/kg; Sigma) following overnight fasting, and continued on HFD until sample collection. Male mice were used due to their greater susceptibility to developing T2DM under this protocol. Control (Ctrl) male mice of matched age, fed a standard laboratory chow diet. Fasting blood glucose was tested one-week post-STZ administration, and mice with glucose levels <11.1 mmol/L were excluded from further analysis.

Postn-CreERT2 transgenic mice were kindly provided by Dr. Xianglong Han. All transgenic mice, including Postn-CreERT2 and tdTomato (Jackson Laboratory, #007909), were maintained on a C57BL/6 background. To induce Cre-mediated recombination, tamoxifen (70 mg/kg) was administered via intraperitoneal injection. Mice received an intraperitoneal injection of EdU (10 mg/kg) 4 hours prior to tissue harvesting, and EdU-positive cells were detected using the Click-iT™ EdU Cell Proliferation Kit (Invitrogen, C10337).

### Cell Isolation and Sequencing

To obtain periosteal cells, femurs and tibias were carefully stripped of muscle, and the bone marrow was removed by flushing with culture medium to minimize hematopoietic contamination. Bones were then digested in collagenase type I (MilliporeSigma). Post-digestion, the cell suspension was centrifuged to pellet the cells, after which the precipitate was resuspended in fresh medium.

For single-cell transcriptomic profiling, femoral and tibial periosteal tissues were collected from control and T2DM mice, and pooled to generate two representative samples for each condition (n = 5 mice/group). scRNA-seq was conducted using the 10x Genomics platform. Single-cell suspensions were captured using a microfluidic platform, where individual cells were encapsulated with barcoded beads. mRNAs from each cell were reverse-transcribed into cDNA. Raw sequencing reads were generated by Illumina NovaSeq 6000 platform and mapped to the mice reference genome (mm10) through Cell Ranger (v7.0). Unique molecular identifiers (UMIs) were subsequently quantified for downstream.

Data were processed in Seurat (v5.1.0). Cells expressing fewer than 500 or more than 6,000 genes, number of genes per UMI for each cell above 0.82, as well as those with mitochondrial gene content exceeding 10% or hemoglobin gene content over 1%, were excluded from downstream analysis. After filtering, 14,627 high-quality cells were retained in the control group and 13,950 in the T2DM group, with a mean sequencing depth of 22,950 reads/cell and 24,643 reads/cell, respectively. Data were normalized using SCTransform (66), regressing out potential confounders including mitochondrial and hemoglobin gene content, as well as cell cycle scores. Batch correction and data integration across samples were conducted using canonical correlation analysis via the “IntegrateLayers” function with default parameters. Clustering was performed using the top 30 principal components (resolution = 0.3), and the cells were visualized with t-distributed stochastic neighbor embedding (t-SNE). Cluster-specific marker genes of clusters and differential gene expression were identified using “FindMarkers,”. CellChat (v2.1.2) (67) was used to infer intercellular communication based on known ligand-receptor pairs.

Total RNA was isolated from THBS1-treated or Ctrl periosteal cells with TRIzol reagent. RNA libraries were prepared with the VAHTS Universal V6 RNA-seq Library Prep Kit and subjected to paired-end 150 bp sequencing on the DNBSEQ-T7 system. Raw reads were aligned to the GRCm38/mm10 genome using HTseq, and transcript abundance was quantified with featureCounts. Differential gene was filtered through DESeq2 (68), applying p < 0.05 and |log₂(fold change)| > 1 as significance thresholds. Functional enrichment analyses (GO and KEGG) were performed with clusterProfiler (v3.18.0) (69).

### Fracture Surgery and microCT Analysis

A skin incision was carefully created along the longitudinal axis of the femoral shaft to establish the femoral fracture model. The mid-diaphysis of the femur was transected using surgical scissors. The midshaft of the femur was transected using scissors. A 27-gauge needle was retrogradely inserted through the knee joint into the femoral canal to serve as an intramedullary fixation pin. The muscle and skin were sutured sequentially using 4/0 gauge sutures. Thrombospondin-1 blocking antibody (15 μl of a 250 μg/ml solution diluted in sterile PBS, Thermo Fisher, clone A6.1) or CD47 blocking antibody (15μg, BioXcell, BE0283) was injected into the fracture site of diabetic C57BL/6J mice. The same concentration of IgG control was injected into the fracture site of the control group to control for damage caused by needle injury. The antibody regimen consisted of three injections, one day apart.

Following euthanasia, femurs were collected and scanned using microCT system (Scanco Medical AG) at 55 kV and 0.145 mA, with a voxel size of 10 μm, enabling clear distinction of mineralized bone from soft tissue and air. To evaluate callus parameters, the fracture callus region was manually contoured using Scanco software, and three-dimensional reconstructions were generated.

### Histology and Immunofluorescence Staining

Mice femurs were carefully dissected to remove surrounding skin and muscle, followed by fixation in 4% paraformaldehyde overnight at 4°C. Tissues were decalcified in 14% Ethylenediaminetetraacetic acid at room temperature for 7 days. After decalcification, femurs were dehydrated in 30% sucrose overnight.

Sections were incubated in 0.5% Triton X-100 in PBS for 10 minutes to permeabilize, followed by PBS washing. Subsequently, tissues were blocked by serum (Invitrogen) for 30 minutes at room temperature. Primary antibodies, including anti-CTSK (1:100; Proteintech, 11239-1-AP), anti-Ki67 (1:100; CST, #9129), anti-COL1A1 (1:50; Beyotime, AF6524), anti-F4/80 (1:100; Abcam, ab300421), and THBS1 antibody (1:100; A6.1, Thermo Fisher), were applied and incubated overnight at 4°C. After washing, sections were incubated with Alexa Fluor 488 (1:500; Thermo Fisher, ab150113) or Alexa Fluor 647 (1:500; Abcam, A21245), for 1 hour at room temperature. For TUNEL staining, the One-step TUNEL FITC Apoptosis Detection Kit (APExBIO, K1133) was used according to the manufacturer’s instructions. Slides were stained with DAPI for nuclear visualization, mounted with anti-fade medium (Invitrogen), and imaged by Olympus confocal microscopy. Image analysis and quantification were completed using ImageJ.

For histological analysis, tissues were immersed in 4% Paraformaldehyde overnight at room temperature, followed by decalcification in 14% Ethylenediaminetetraacetic acid for 2 weeks. Masson’s trichrome staining (Solarbio, G1340) and Safranin O/Fast Green staining (Solarbio, G1371) were performed following the manufacturer’s protocols for brightfield imaging.

### Mitochondrial Function Assays

For mitochondrial assays, cells were incubated separately with either 500 nM MitoSOX™ Red (Thermo Fisher, M36007) or 100 nM TMRE (Thermo Fisher, T669), each in combination with 100 nM MitoTracker™ Green FM (Thermo Fisher, M7514) and Hoechst 33342 (Thermo Fisher, H3570) for 30 minutes at 37°C. After washing, cells were maintained in fresh medium and imaged using an Olympus FV3000 confocal microscope to assess mitochondrial ROS, membrane potential, and morphology.

### Western Blot

Proteins were analyzed by Western blot using anti-LaminB1 (1:1000; Beyotime, AF1408) and anti-β-Actin (1:1000; Beyotime, AF5003) antibodies. The PVDF was incubated with goat anti-rabbit IgG (1:5000, SAB, L3012) and visualized using an ECL kit (Vazyme, E412-01).

## Statistics

Data were presented as mean ± standard deviation (S.D.). Statistical analysis was performed using Prism 9 software. For comparisons between two groups, an unpaired Student’s t-test was used, whereas multiple group comparisons were analyzed by one-way ANOVA followed by Tukey’s multiple comparisons test.

## Acknowledgments

**Funding:** This work was supported by the Sichuan Science and Technology Program (Grant No: 2025ZNSFSC0054 to Y.S.); Sichuan Science and Technology Program (No.24ZDYF0099 to J.W.); Research and Develop Program, West China Hospital of Stomatology Sichuan University (Grant No: RD-03-202304 to Y.S.); Funding of the State Key Laboratory of Oral Diseases (Grant No. SKLOD-2025KP005 to Y.S.).

## Author contributions

Conceptualization, Fangyuan Shen, Ling Ye and Yu Shi. Software, Fangyuan Shen. Validation, Fangyuan Shen, Moyu Liu, Qiaoyue Ren. Methodology, Fangyuan Shen, Moyu Liu, Qiaoyue Ren, and Ting Zheng. Writing, Fangyuan Shen, Yu Shi and Ling Ye. Resources, Fangyuan Shen, Moyu Liu, Qiaoyue Ren, Ting Zheng. Data Curation, Fangyuan Shen, Moyu Liu, Qiaoyue Ren, Jingyao Cui, and Puying Yang. Fund support: Jun Wang and Yu Shi. Fangyuan Shen and Yu Shi were responsible for the decision to submit the manuscript.

## Data and materials availability

The authors confirm that the data supporting the findings of the present study are available within the article and its Supporting Information. Raw data of RNA-seq and scRNA-seq have been deposited in the Gene Expression Omnibus repository and can be accessed via GSE294691 and GSE294693.

