## Supplementary material for "Immune-Derived THBS1-CD47 Axis Induces Cellular Senescence and Suppresses Osteogenesis in Diabetic Periosteum": Figs. S1 to S5 & Materials and Methods

Fangyuan Shen *et al.*

**This PDF file includes:**

Figs. S1 to S5

Materials and Methods

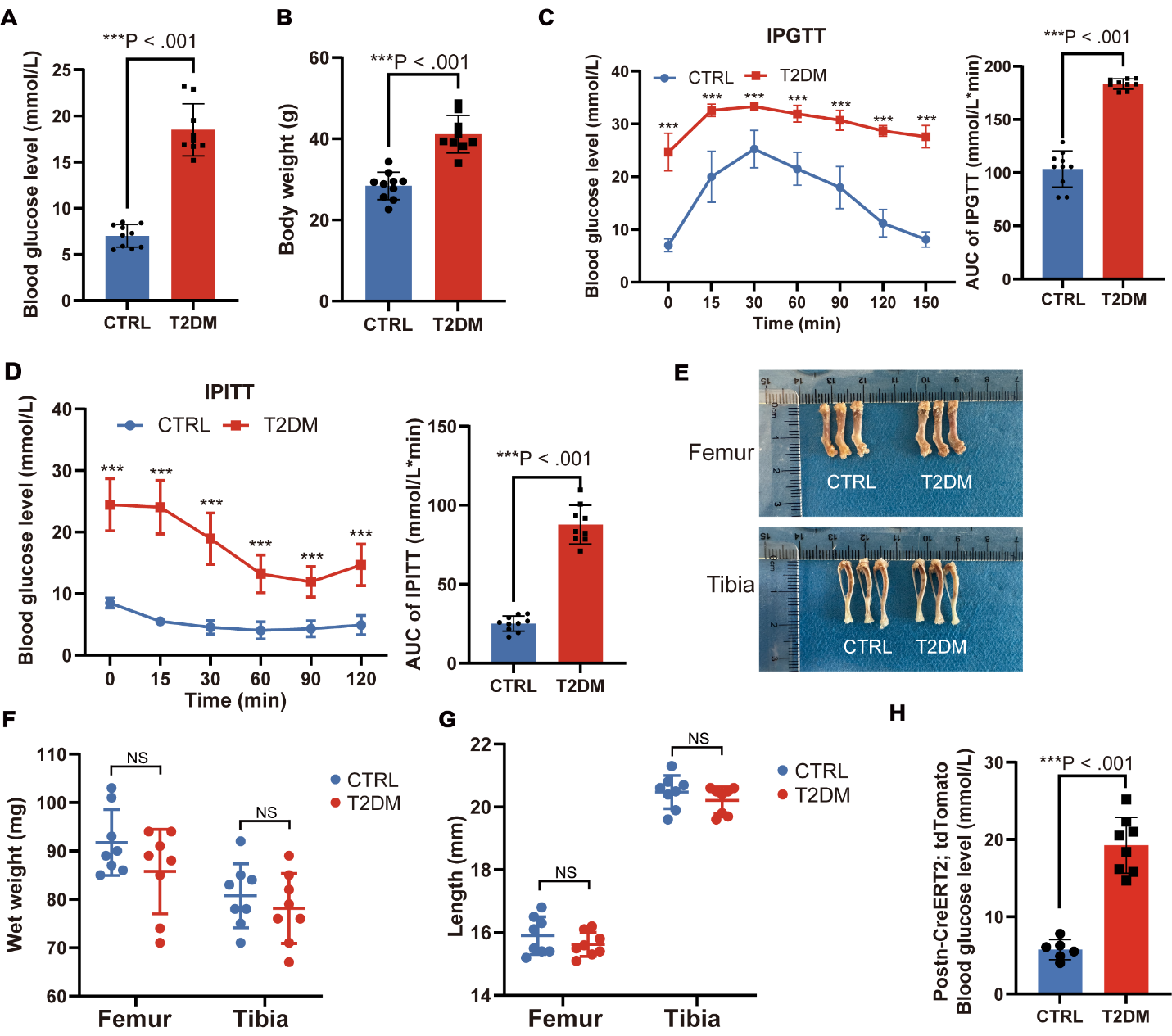

Fig. S1.

(**A**) Blood glucose levels in C57BL/6J mice after 8 hours of fasting. (**B**) Body weight of T2DM and control mice before sacrifice. (**C** and **D**) Intraperitoneal glucose tolerance test (IPGTT) glucose curve and Intraperitoneal Insulin tolerance test (IPITT) glucose curve, followed by calculating the integrated area under the curves (AUC). CTRL, n=10; T2DM, n=9. (**E**) Representative gross images of femur and tibia from CTRL and T2DM groups. (**F**) The graph was the quantitative Wet weight of femur and tibia (n = 8 per group). (**G**) The graph showed length of femur and tibia (n = 8 per group). (**H**) Blood glucose levels in Postn-CreERT2; tdTomato mice after 8 hours of fasting. CTRL, n=3; T2DM, n=4. All quantitative data are presented as mean ± S.D. Each data point represents one animal (biological replicate). Statistical analysis was performed using an unpaired Student’s t-test.

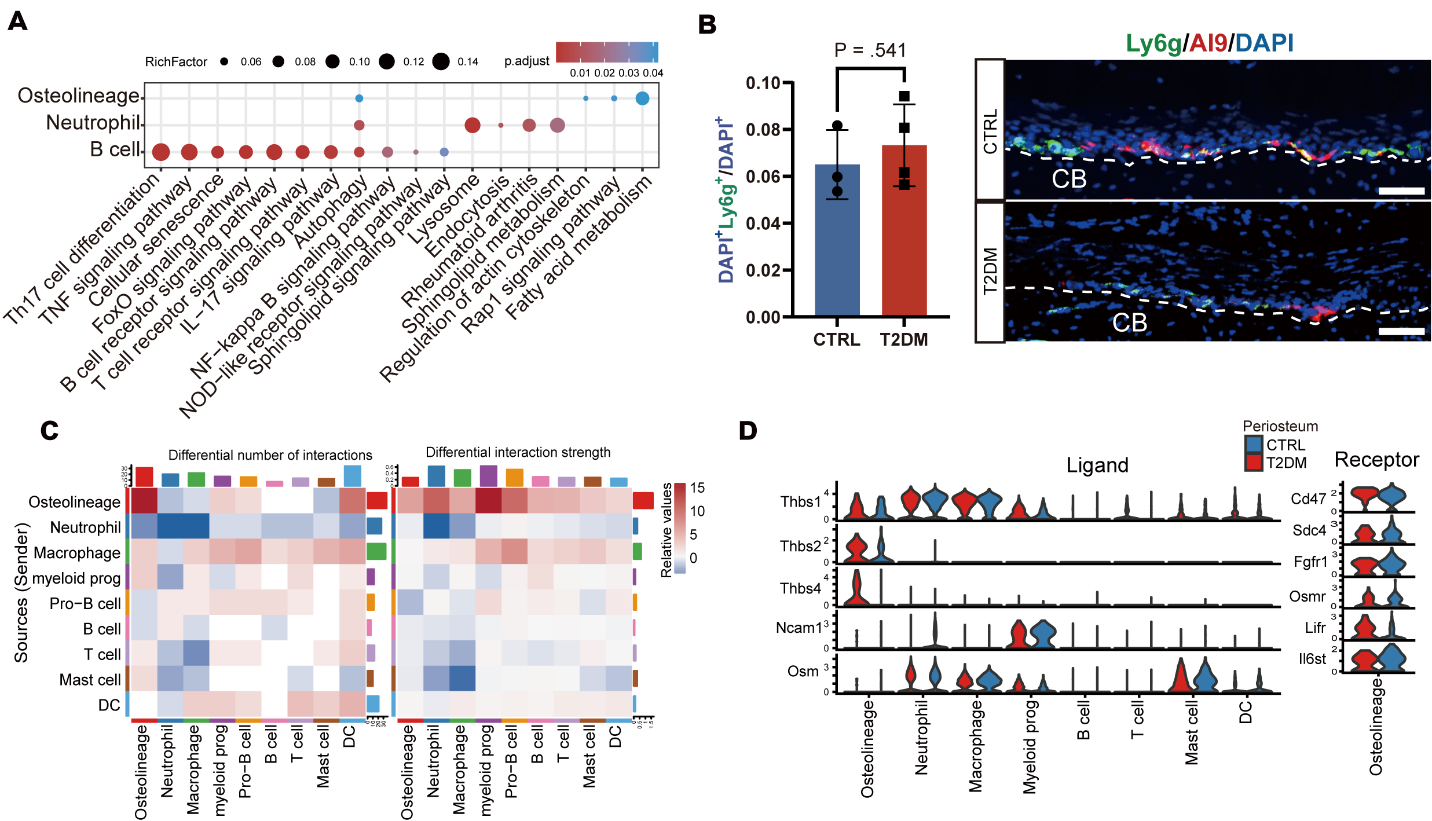

Fig. S2.

(**A**) KEGG pathway enrichment analysis of upregulated genes in osteolineage, neutrophil, and B cell. (**B**) Immunofluorescence staining and quantification of Ly6g in the periosteum of T2DM (n=4) and CTRL (n=3) groups. Quantitative data are presented as mean ± S.D, Scale bar: 100 μm. Each data point represents one animal (biological replicate). Statistical analysis was performed using unpaired Student’s t-test. (**C**) Differential cell communication statistics of each cluster. (**D**) Periosteal expression levels of each ligand and receptor involved in the differential ligand-receptor interactions.

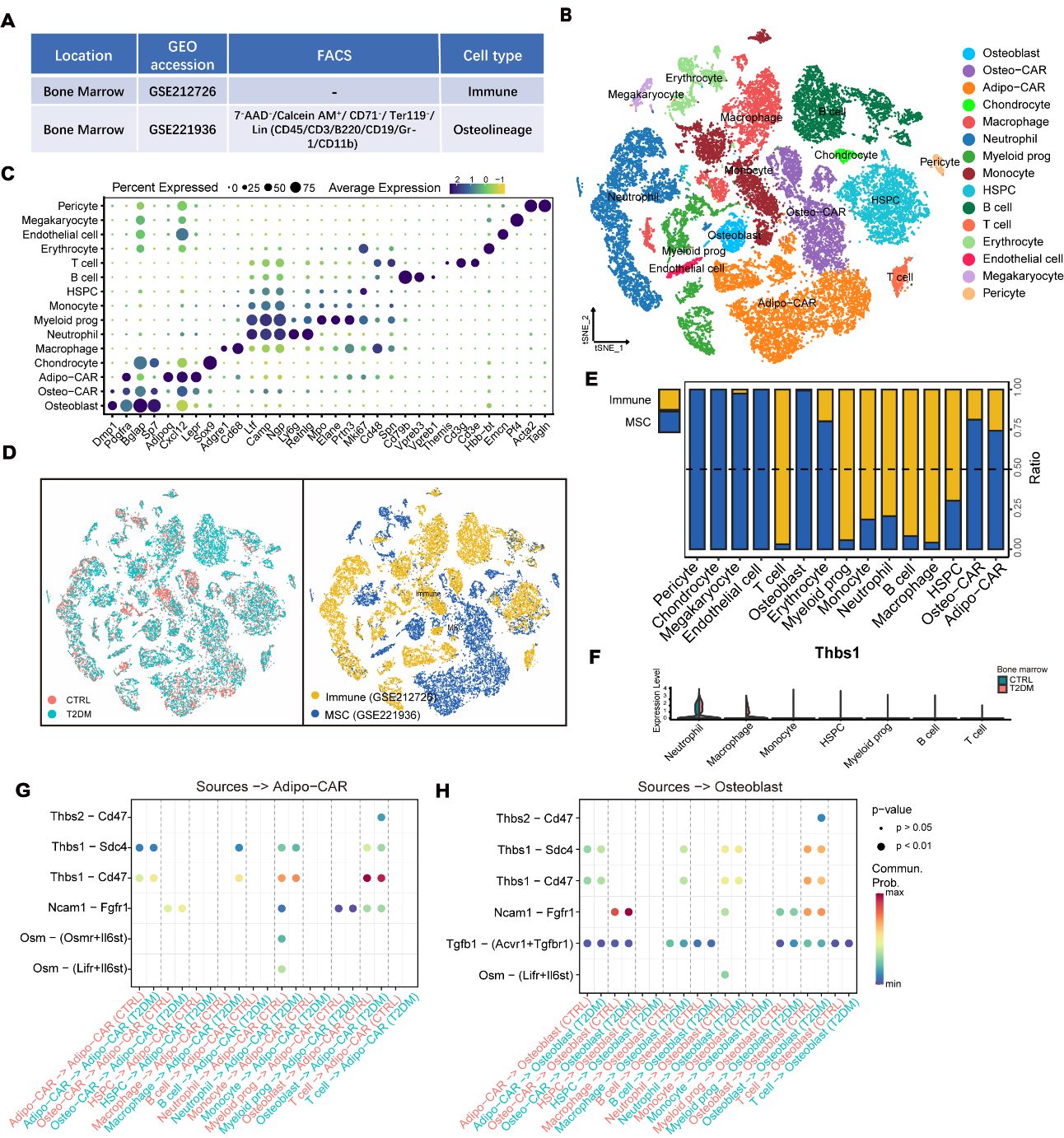

Fig. S3.

(**A**) T2DM and CTRL mouse bone marrow single-cell sequencing data information. (**B**) tSNE plot from scRNA-seq of integrated bone marrow immune cells and osteolineage cells from T2DM and CTRL mice. (**C**) Dot plots of marker genes are used to annotate each identified cluster. (**D**) tSNE diagram showing the distribution of T2DM and CTRL as well as immune cells and MSCs. (**e**) Bar graph showing the relative proportions of each cell cluster in immune cell and MSC. (**F**) Expression levels of Thbs1 in immune cells of bone marrow. (**G** and **H**) Differential ligand-receptor interactions from immune cells to adipo-CAR and osteoblast in the bone marrow of T2DM and CTRL group.

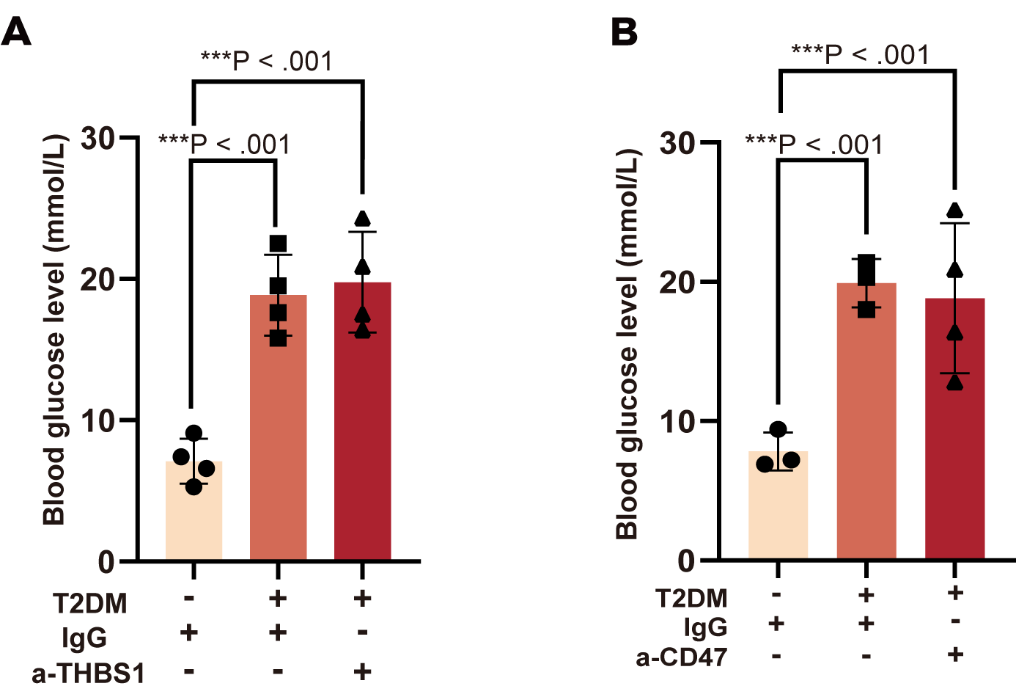

Fig. S4.

(**A-B**)Blood glucose levels of fracture mice in each group after 8 hours of fasting before surgery. Data are presented as mean ± S.D. At least three biological replicates were analyzed. Statistical significance was determined using one-way ANOVA followed by Tukey’s multiple comparisons test.

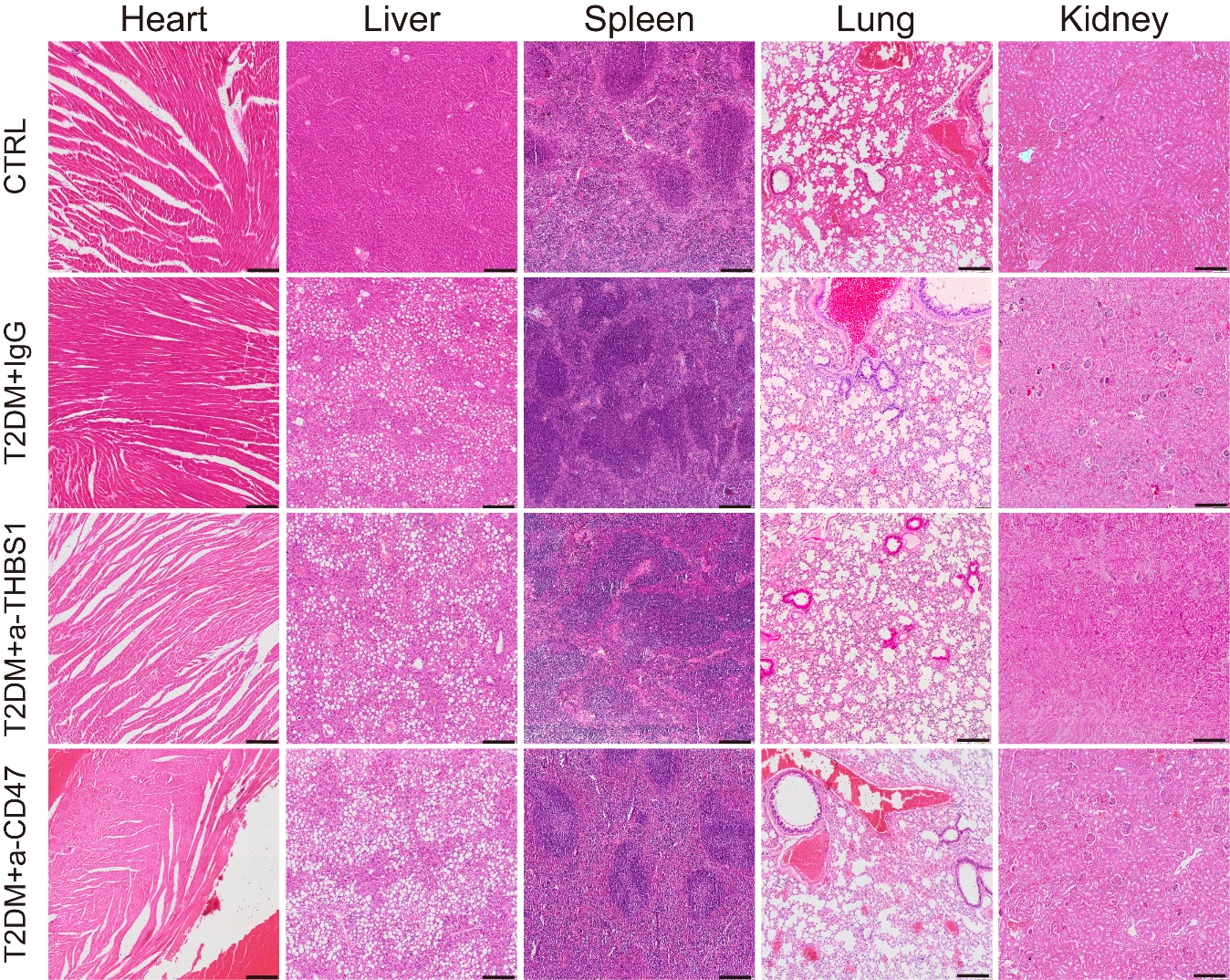

Fig. S5.

H&E staining of important organs in different treated mice. Scale bars: 200 μm.

Materials and Methods

**Seahorse assay**

Oxygen consumption rate (OCR) was measured using the Seahorse XFe24 Analyzer (Agilent) following the manufacturer’s protocol. Periosteal cells were seeded at 40,000 cells per well in Seahorse XF24 microplates. After 2 hours, the medium was replaced with Seahorse XF assay medium containing 5.5 mM glucose, 2 mM glutamine, 0.1 mM pyruvate, and 5 mM HEPES. For the Mito Stress Test, 1.5 μM oligomycin, 1.5 μM FCCP, 1 μM Rotenone, and 1 μM Antimycin A were sequentially injected. Data were analyzed using Seahorse Wave software.

**qPCR**

Total RNA was extracted using TRIzol, reverse-transcribed with a Vazyme RT kit (R223-01), and analyzed by qPCR using ChamQ™ Universal SYBR® qPCR Master Mix (Vazyme, China).

**Flow cytometry and cell sorting**

Periosteal and bone marrow cells were freshly isolated from the femur and tibia of control and T2DM mice, followed by red blood cell lysis and filtration through a 70-μm strainer to obtain single-cell suspensions. Cells were stained with a panel of fluorochrome-conjugated antibodies, including PerCP-Cy5.5-7-AAD (Invitrogen, 6993-50), APC-CD47 (BioLegend,127513), APC-F4/80 (BioLegend,123115), PE-Cy7-CD45 (BioLegend, 101207), FITC-Ly6G (BioLegend, 164507), and PE-CD11b (BioLegend, 101207). After incubation on ice for 30 min in the dark, cells were washed twice with cold PBS containing 2% FBS and resuspended for analysis. Flow cytometry was performed CytoFLEX and cell sorting was conducted using a BD FACSAria™ III. Data were analyzed using FlowJo software (TreeStar). Gating strategies were applied to exclude debris and doublets, and live CD45⁺CD11b⁺F4/80⁺ macrophages and CD45⁺CD11b⁺Ly6G⁺ neutrophils were sorted separately from periosteum and bone marrow for downstream qPCR analysis.
